# Auxin requirements for a meristematic state in roots depend on a dual brassinosteroid function

**DOI:** 10.1101/2020.11.24.395483

**Authors:** M. Ackerman-Lavert, Y. Fridman, R Matosevich, H Khandal, L. Friedlander, K. Vragović, R. Ben El, G. Horev, I Efroni, S. Savaldi-Goldstein

**Affiliations:** Faculty of Biology, Technion-Israel Institute of Technology, Haifa 3200003, Israel; Institute of Plant Sciences and Genetics in Agriculture, Faculty of Agriculture, The Hebrew University of Jerusalem, Rehovot, Israel; Lorey I. Lokey Interdisciplinary Center for Life Sciences and Engineering, Technion–Israel Institute of Technology, Haifa 3200003, Israel

## Abstract

The organization of the root meristem is maintained by a complex interplay between plant hormones signaling pathways that both interpret and determine their accumulation and distribution. Brassinosteroids (BR) and auxin signaling pathways control the number of meristematic cells in the Arabidopsis root, via an interaction that appears to involve contradicting molecular outcomes, with BR promoting auxin signaling input but also repressing its output. However, whether this seemingly incoherent effect is significant for meristem function is unclear. Here, we established that a dual effect of BR on auxin, with BR simultaneously promoting auxin biosynthesis and repressing auxin transcriptional output, is essential for meristem maintenance. Blocking BR-induced auxin synthesis resulted in rapid BR-mediated meristem loss. Conversely, plants with reduced BR levels were resistant to loss of auxin biosynthesis and these meristems maintained their normal morphology despite a 10-fold decrease in auxin levels. In agreement, injured root meristems which rely solely on local auxin synthesis, regenerated when both auxin and BR synthesis were inhibited. Use of BIN2 as a tool to selectively inhibit BR signaling, revealed meristems with distinct phenotypes depending on the perturbed tissue; meristem reminiscent of BR-deficient mutants or of high BR exposure. This enabled mapping BR-auxin interactions to the outer epidermis and lateral root cap tissues, and demonstrated the essentiality of BR signaling in these tissues for meristem maintenance. BR activity in internal tissues however, proved necessary to control BR homeostasis. Together, we demonstrate a basis for inter-tissue coordination and how a critical ratio between these hormones determines the meristematic state.

## Introduction

Throughout the plant’s life span, plant meristems supply cells to the developing above- and below-ground organs. From a developmental point of view, auxin is essential for the establishment and maintenance of a functional root meristem; a severe drop in auxin levels, as seen in auxin biosynthesis mutants, leads to meristem loss [1–4]. The production of endogenous auxin indole-3-acetic acid (IAA) occurs in several tissues of the root meristem, but it is the specific production in the quiescent center (QC) cells and its distribution by the polar auxin transport machinery, that form auxin maxima critical to maintain the meristem [2, 5, 6]. IAA is produced from L-tryptophan (Trp) in a two-step process involving the tryptophan aminotransferases family (e.g. TRYPTOPHAN AMINOTRANSFERASE OF ARABIDOPSIS1 (TAA1) and the flavin monooxygenases YUCCA (YUC) family [7].

BR regulates the size of the meristem [8, 9]. Low BR levels attenuate cell cycle activity while supra-optimal levels, as achieved upon exogenous application of BR, limit meristematic cell counts, promoting their differentiation. The reported complex molecular interplay between BR and auxin in the root meristem has both positive and negative aspects [10]. Within a positive interaction, BR elevates the transcription of select auxin biosynthesis genes, e.g., YUC and TAA1 in the outer tissues, as concluded from a translatome mapping of the BR response [11]. High BR levels also elevate auxin signaling input in the nucleus of meristematic cells by repressing PIN-LIKES (PILS), a putative intracellular auxin transport facilitator [12]. BR was also observed to relieve the low nuclear auxin input in root meristem of phloem development mutants [13]. Within a negative interaction, BR and auxin have opposing effects on the transcription of common target genes [14]. The negative molecular interaction between BR and auxin is implicated within their antagonistic effect on QC divisions [9, 14], and within root elongation assays [15–17]. However, elevation of nuclear auxin input, as found in select *pils* mutants, did not counteract the BR effect on the meristem [12]. Together, whether this seemingly incoherent molecular interaction between the two hormones indicates a dual BR effect on auxin and the relevance of such a putative effect to meristem function, remain unclear.

BR signaling in roots is tissue- and zone-dependent [8, 11, 18–20]. It is relatively low in the meristem and increases in the basal meristem, forming a gradient in the elongation zone, particularly in the epidermis [14]. This BR signaling pattern contrasts the high auxin signaling in the maxima zone [14], but is similar to the auxin signaling gradient in the epidermis [10]. In addition, the epidermis has highest gene response to BR as compared to inner tissues [11].

Limiting the expression of the broadly expressed BR receptor BRI1 to the epidermis, elevated auxin biosynthesis genes, promoted cell cycle activity and delayed cell differentiation accompanied by an extended zone of auxin maxima [8, 11]. BRI1 in the inner stele tissue, however, counteracts and thus balances this epidermal effect [11]. The relevance of the reported positive and negative nature of BR-auxin interactions to meristem maintenance and how they are integrated with the spatiotemporal effect of BR signaling remain unclear.

In the canonical BR signaling pathway, BRI1 perceives BR with its co-receptor BAK1 and the signal is then transduced by several well established regulatory steps involving several proteins. Among these is the GSK3 kinase BIN2, which plays a major inhibitory role by phosphorylating and thus inhibiting the activity of key downstream transcription factors belonging to the BES1/BZR1 family, reviewed in [21, 22]. High BR levels relieve the inhibition of BIN2, enabling the accumulation of BES1/BZR1 in the nucleus, where they activate and repress many genes [23–25]. Among these genes, select BR biosynthesis genes are repressed, driving a negative feedback mechanism that restrains BR signaling output [26–28]. Active BRs are derived from multiple biosynthesis steps acting in parallel routes [29–31]. Since the discovery of the BR biosynthesis genes more than three decades ago, clear-cut evidence of which root tissue produce BRs is still lacking. Root tissue-specific transcriptional data sets suggest that BR biosynthesis genes have diverse expression patterns [11, 31], and can be repressed in response to BR in various tissues [11]. However, the importance of this feedback in meristem function and how it integrates with current knowledge of spatiotemporal BR signaling, remains uncertain.

Here, we expand the experimental “BR toolkit” by establishing lines with tissue-specific expression of BIN2, a protein that selectively inhibits BR signaling. By combining a pharmacological approach, usage of available mutants and auxin markers and direct hormone measurement, we revealed the role of the dual effect of BR on auxin and mapped the essentiality of BR for this interaction and for meristem homeostasis.

## Results

### BR promotes auxin synthesis and levels but simultaneously inhibits auxin signaling output

To determine the association between auxin levels and auxin signaling in response to BR, we quantified the levels of YUC9 and the auxin signaling input- and output, in brassinolide (BL) treated plants along the epidermis. First, *pYUC9-VENUS-YUC9ter* plants were subjected to moderate BL treatment (0.8 nM) for 12h (Figure 1A,B). A pronounced elevation of the fluorescent signal was observed in the root epidermis (Figure 1A,B). Moreover, a direct measurement of IAA levels revealed that BL treatment increased auxin levels in the root tip (Figure 1C). In addition, the signal corresponding to the *DII-venus* auxin sensor [32] was lower in response to this same BL treatment, indicating high auxin signaling input (Figure 1D, E). Thus, auxin signaling input is increased by exogenous moderate BR, as was seen following short-term exposure to high BR levels [12]. To determine whether this enhancement in auxin input was an outcome of BR-mediated elevation of auxin biosynthesis, seedlings were exposed to BL in the presence and absence of L-Kynurenine (Kyn) that moderately inhibits auxin biosynthesis [33]. As expected, addition of Kyn enhanced the DII-venus signal intensity (Figure 1D,E). In contrast, BL in the presence of Kyn reduced the signal with a relatively lower fold change as compared to BL alone, indicating that the BL effect largely depends on IAA production. Conversely, blocking BR biosynthesis by brassinazole (BRZ) [34] for the same duration, increased the signal intensity of the sensor (Figure 1D,E). Taken together, BR elevates auxin biosynthesis in the epidermis and in accordance, auxin levels and thus auxin signaling input.

**Figure 1.**
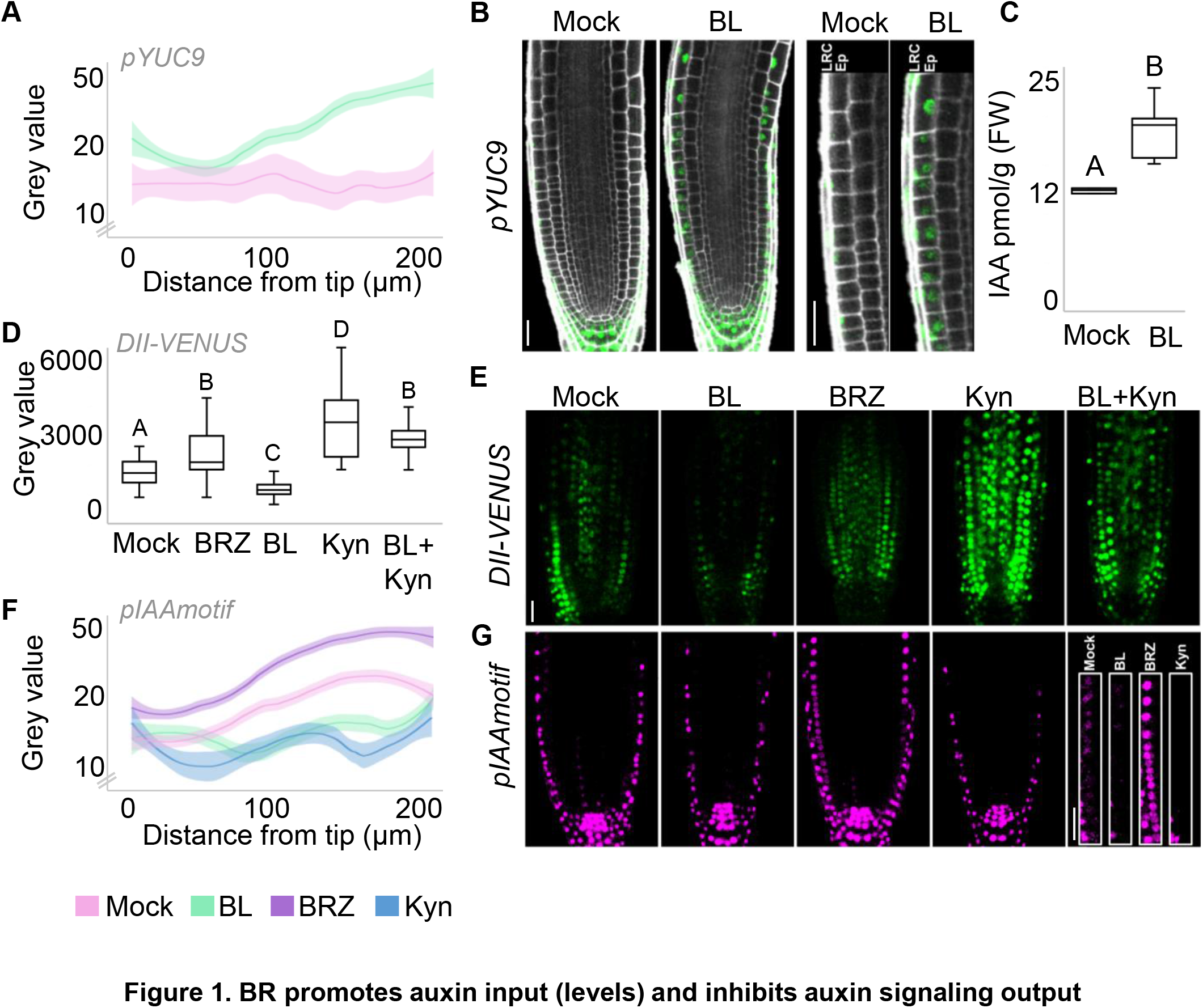
BR promotes auxin input (levels) and inhibits auxin signaling output. (A,B) *pYUC9-VENUS-termYUC9* roots treated with DMSO (mock) or 0.8nM brassinoslide (BL), for 12h. (A) Quantification of the epidermal signal (log10 scale) along the meristem (n = 13 and 17 for mock and 0.8 nM BL respectively), captured by confocal microscopy. (B) Confocal images showing fluorescent signals of VENUS (green) and propidium iodide (PI, in white). Right panels show a close-up of the corresponding roots, to highlight BL-mediated elevation of YUC9 in the epidermis (ep). Lateral root cap (LRC). (C) IAA content in root tips treated with mock and 0.8 nM BL for 12 h. Different letters indicate values with statistically significant differences (p ≤ 0.05, according to Tukey HSD). (n=5) (D,E) Roots expressing *DII-VENUS* treated with mock, 0.8 nM BL, 3 μM BRZ, 4 μM Kyn or BL+Kyn, for 12h. (D) Quantifications of the *DII-VENUS* fluorescent signal in the meristem, as captured using confocal microscopy. (E) Representative confocal images corresponding to the quantification in (D). Different letters indicate values with statistically significant differences (p ≤ 0.05, according to Tukey HSD; n = 49, 27, 39, 40 and 29 for mock, 0.8 nM BL, 3 μM BRZ, 4 μM Kyn and BL+Kyn respectively). (F,G) *plAAmotif-mScarlet-NLS* roots treated with mock, 0.8 nM BL, 3 μM BRZ or 4 μM Kyn, for 12h. (F) Quantification of the fluorescence signal (log_10_ scale) in the epidermis along the meristem (n = 20, 27,15 and 14 for mock, 3 μM BRZ, 0.8 nM BL, and 4 μM Kyn respectively). (G) Confocal images showing the *mScarlet* fluorescence signal (magenta). Right panels show a close-up of the corresponding roots to highlight low and high fluorescence signals in response to BL and BRZ, respectively. Scale bars = 25 μm.

Next, we quantified the auxin signaling output. BR treatment of the double reporter *DR5v2-ntdTomato–DR5-n3GFP* [35] led to downregulation of the epidermal DR5v2-ntdTomato signal (Figure S1), while BRZ treatment elevated it. The DR5-n3GFP signal, primarily in the QC and surrounding cells, was mildly increased or remained unaffected (data not shown). In addition, the sensitive output readout pIAAmotif, that is also expressed in the epidermis [36] revealed that BL reduced the signal measured in the epidermis as with the addition of Kyn, while BRZ treatment elevated it (Figure 1F,G). A similar trend was observed in the stele (Figure S1). In conclusion, BR promotes auxin levels and thus signaling input but simultaneously inhibits auxin signaling output in the same tissue.

### BR-mediated auxin elevation is critical for meristem maintenance

To evaluate the impact of this dual effect of BR on meristematic function, mutants in auxin biosynthesis genes *wei2,wei7* (hereafter called *wei2,7*) and *yuc3yuc8yuc9yuc5* (hereafter called *yuc-q*) [3, 37] were subjected to the moderate BL treatment. In the absence of BL treatment, the roots of *wei2,7* have an overall normal meristem structure, but with reduced numbers of meristematic cells and shorter roots than wild type plants (Figure S2). Remarkably, while the meristematic cell counts in wild-type plants were slightly reduced by the BL treatment, the meristem of *wei2,7* was completely lost within 96h (Figure 2A). Addition of exogenous auxin (NAA) restored the mutant meristem in the presence of BL (Figure 2A). A similar BR-mediated meristem loss was observed in *yuc-q* (Figure 2A). To test whether endogenous production of auxin in the epidermis is sufficient to counteract the differentiation effect of BL, the YUC5 was driven by the *pGL2* promoter in wild type and in a *yuc-q* mutant background. Roots of *pGL2-YUC5* were short and hairy, while the root meristem had more cells, all indicative of a high auxin phenotype (Figure S2). Importantly, *pGL2-YUC5;yuc-q* maintained an intact meristem in response to BL (Figure 2A).

**Figure 2.**
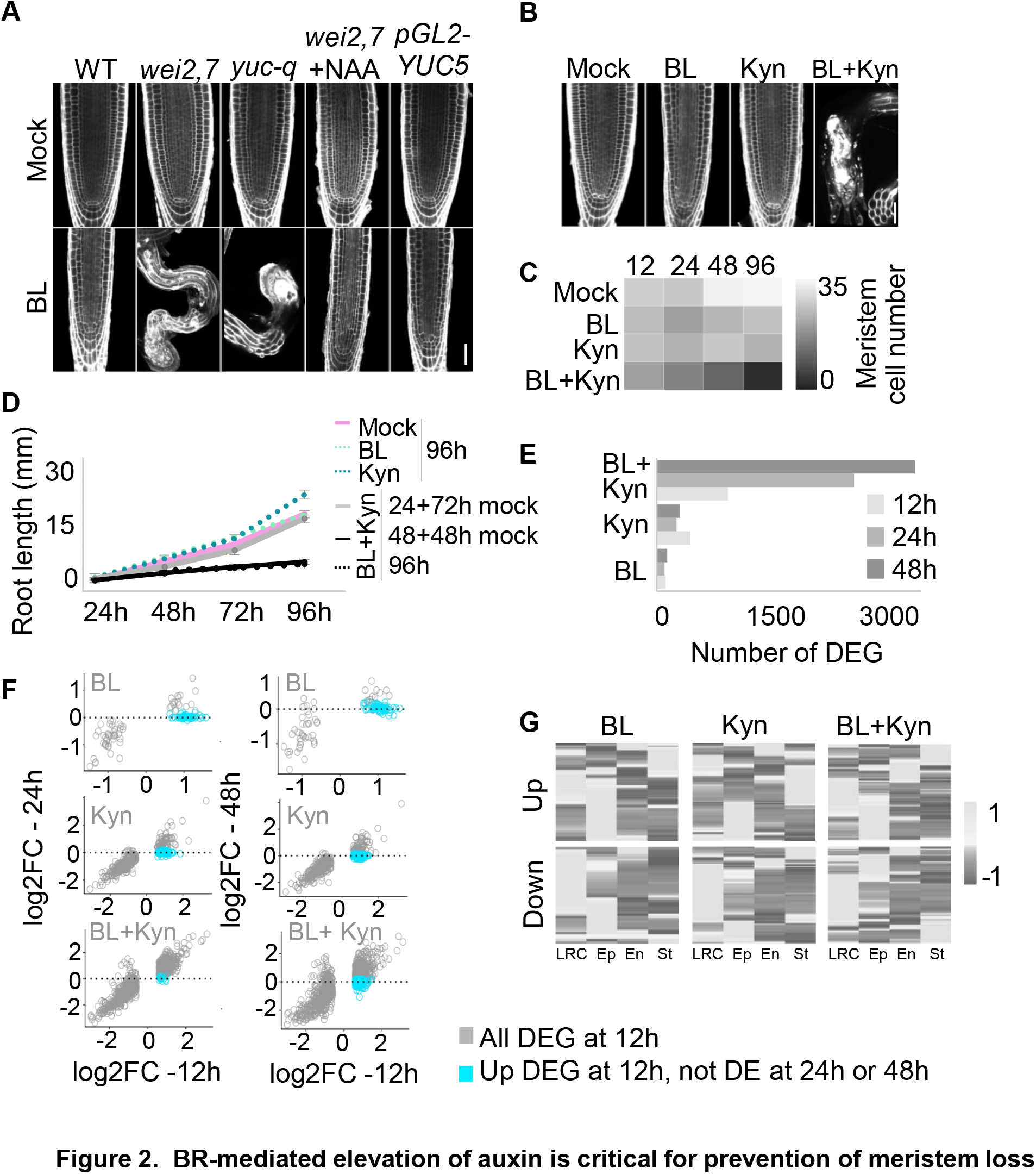
BR-mediated elevation of auxin is critical for prevention of meristem loss. (A, B) Confocal microscopy images of representative root meristems 96h after transfer to the indicated treatment. (A) wild type (WT), *wei2,7, yuc-q* and *yuc-q* expressing *pGL2-YUC5* after transfer to mock or 0.8 nM BL*. wei2,7* were also transferred to NAA (0.2μM) (see also Figure S2). (B) Wild-type meristems after transfer to mock, 0.8 nM BL, 4 μM Kyn or their combinations. PI fluorescence signal in white (see also Figure S3). (C) Number of meristematic cells in roots subjected to hormonal treatments as in (B) for 12, 24, 48 or 96 h, presented as a heatmap (n ≥10, see also Figure S3). (D) Root length of seedlings exposed to mock, BL, Kyn and the combined BL+Kyn treatments as in (B) for a limited duration of time (24, 48 or 96h), were transferred to mock conditions for recovery (n≥20). (E) Number of differentially expressed genes (DEGs) in response to mock, 0.8 nM BL, 4μM Kyn and their combinations after 12h, 24h and 48h (see also Figure S3 and Table S1). (F) Scatterplots comparing DEGs at 12 h (x-axis, gray dots) with the same genes at 24 h (left panels, y-axis) and at 48 h (right panels, y-axis) after each treatment. DEGs at 12 h treatment (p≤0.05, and FC≥ 1.5) that are not differently expressed on the y-axis (24h or 48h) are marked in blue (see also Figure S4). (G) Expression of DEGs at 12 h as in (E) in different root tissues (epidermis, LR/C, endodermis and stele) (see also Figure S4). Scale bars =25 μm.

Application of both Kyn and BL caused a rapid reduction of meristematic cell numbers, leaving approximately half of the cells by 12h, until complete meristem loss by 96h. Kyn and BL alone however reduced the number of meristematic cells over time (Figure 2B,C), without causing meristem loss (Figure S3). In addition, the meristem remained intact by a single treatment of 125-fold more BL or 12-fold more Kyn by the 96h tested (Figure S3). When exposing roots to the combined hormonal treatments for a limited duration of time, and then transferring them to mock conditions for recovery (Figure 2D), the meristem and in accordance, root growth, could be recovered. However, after 48h of treatment, the meristem was not recovered and committed to differentiation (Figure 2D). In agreement with the observed meristematic phenotype, RNA-seq analysis showed that only about 100 genes were differentially expressed in response to BL at each tested time point, until 48h (Table S1, Figure 2E). This reflects the moderateness of the perturbation protocol used. Kyn treatment triggered more differentially expressed genes (DEGs) as compared to BR, but as seen with the BR treatment, the number of DEGs did not increase with time (Table S1, Figure 2E). However, the combined treatment had a synergistic effect on the number of DEGs, which was already noted at the first time point analyzed (i.e., 12h). This synergistic effect increased with time, reaching up to 30-fold increase in the number of DEGs as compared to the single-hormone perturbations, showing concomitant repression of meristematic genes and induction of elongation and differentiation zone genes (Table S1, Figure 2E, Figure S4). In response to BL and Kyn alone, the majority of DEGs measured at 12h were back to their basal expression levels by 24h post-treatment (75% for BL and 53% for Kyn, Figure 2F, Figure S4). However, following combined treatment, only 3% of DEGs were back to their basal level after 24h and expression patterns were generally maintained over 48h (Figure S4). This suggests that changes in single hormone levels involve feedback on gene expression that is lost when the dual function of BR is perturbed. Together, BR-mediated activation of auxin synthesis is critical to buffer its opposing effect on auxin and thus to ensure a meristematic state.

### The inevitable meristem loss caused by low auxin is blocked in the absence of BR

If a dual effect of BR is active in the root meristem, then the meristem of BR-deficient mutants would be less affected by a reduction in auxin synthesis. Treatment of wild type seedlings with high Kyn doses (50-200 μM) led to a fully differentiated (or highly deformed) meristem within 10-14 days (Figure 3A,B, Figure S5). Strikingly, *bri1* mutants (two independent loss-of-function alleles tested), *cpd* (a BR biosynthesis mutant) and BRZ-wild type treated seedlings maintained an intact root meristem under these deleterious reduction in auxin synthesis (Figure 3A,B, Figure S5). Direct measurement of IAA levels confirmed that Kyn treatment strongly reduced auxin levels, even in the presence of BRZ (Figure 3C). Interestingly, despite lower auxin levels, the distribution of auxin signaling input appeared normal and auxin transcriptional output was still maintained in the stem cell niche (Figure 3D,E; Figure S5). Finally, treating assessment of *bri1* mutants in combination with mutants in its two homologous receptors *brl1* and *brl3* (i.e., triple mutant) and *bin2-1* (i.e., a dominant version of the inhibitory kinase) validated that BRI1 via inhibition of BIN2, is required for initiation of the differentiation response (Figure S6). Together, low BR levels highly sensitize the root meristem to auxin in the meristem in agreement with a dual BR function effect occurring at a steady state growth. One could also envision however that a generic program could still function in the absence of both hormones.

**Figure 3.**
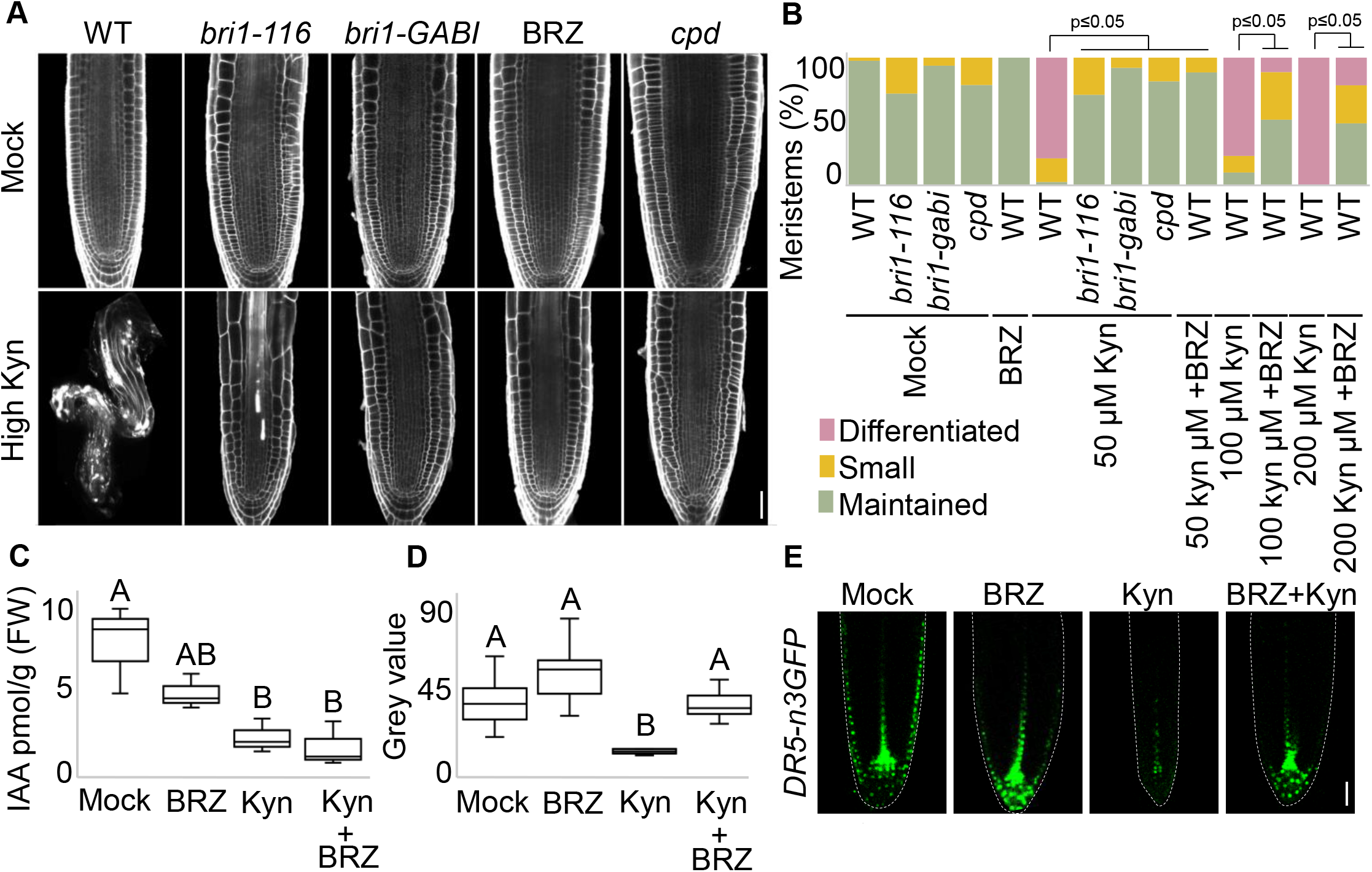
The inevitable meristem loss caused by low auxin is blocked in the absence of BR. (A, B) Confocal microscopy images of representative root meristems of wild-type (WT), WT germinated on BRZ, *bri1-116, bri1-GABI* and *cpd*, 10 days after transfer to mock or high-Kyn (50 μM) conditions (see also Figure S5). (B) Distribution of meristematic phenotypes following long-term high-Kyn treatment, as in (A) (see also Figure S5). For mock treatment, n=83, 32, 16 and 28 for WT, *bri1-116, bri1-GABI* and *cpd*, respectively. For WT+BRZ, n=62. For 50 μM Kyn treatment, n = 113, 24, 25, 27 for WT, *bri1-116, bri1-GABI* and *cpd*, respectively. For WT+BRZ+50 Kyn, n= 78. For 100 μM Kyn treatment, n=39 and for 100 μM Kyn+BRZ treatment, n=35. For 200 μM Kyn treatment, n=49 and for 100 μM Kyn+BRZ treatment n=37. P values are according to Fisher exact test. (C) IAA content in roots treated with mock, high Kyn, BRZ and their combination for 96 h. (D, E) *DR5-n3GFP* roots treated for 13 days (long-term) with mock, BRZ, high Kyn or high Kyn+BRZ (D) Quantifications of the *DR5-n3GFP* fluorescence signal in the QC and surrounding initials, as captured by confocal microscopy, as in (E) (13≤n≤19). (E) *DR5-n3GFP* fluorescence signal (green), dashed lines mark meristem borders (see also Figure S5). Different letters indicate values with statistically significant differences (p ≤ 0.05, according to Tukey HSD). Scale bars =25 μm.

### BR levels dictate auxin-dependent meristem regeneration

Our findings thus far suggest that BR determines the amount of auxin needed to maintain a meristematic state. To determine whether this principle extends beyond steady state, as in the process of meristem regeneration that depends on a critical elevation of auxin after wounding [38], we performed decapitation of the stem cell niche and added Kyn, BL, or BRZ. Addition of BL blocked meristem regeneration and this inhibitory effect of BL was limited to the first 24h after wounding, in agreement with the relatively low auxin levels at the incipient of meristem regeneration (Figure 4A). Indeed, the severity of the inhibitory effect of BL was alleviated with the addition of exogenous auxin, in a concentration-dependent manner (Figure 4B). Meristem regeneration is blocked if auxin synthesis is inhibited, as with the addition of kyn (Figure 4C). Remarkably, lowering endogenous BR levels by BRZ treatment, resisted the deleterious effect of Kyn, allowing meristem regeneration in the presence of low levels of auxin (Figure 4C). Thus, BR dictates auxin requirements for meristematic function within two developmental contexts - steady state growth and regeneration.

**Figure 4.**
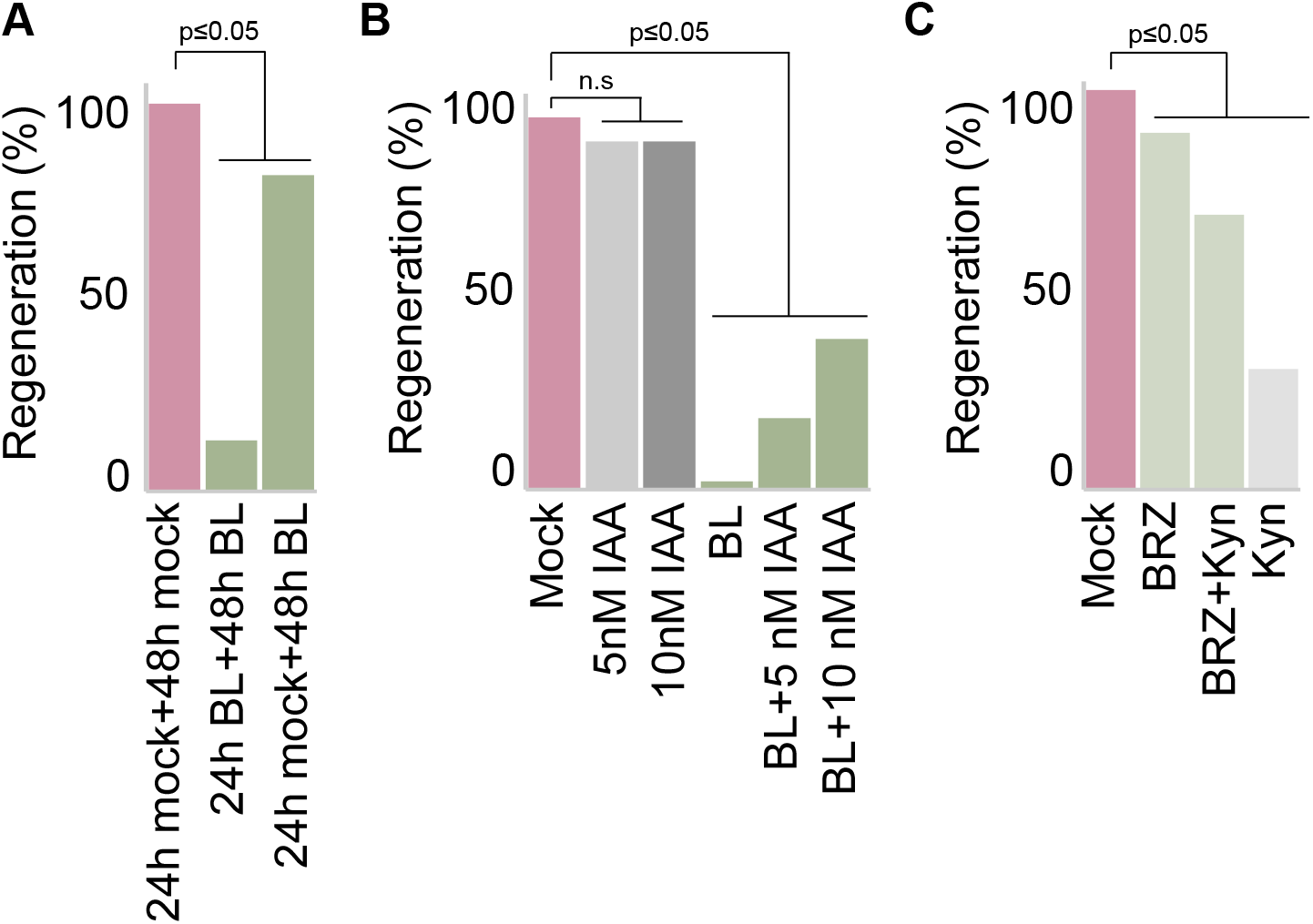
BR levels dictate auxin-dependent meristem regeneration. (A-C) Regeneration rates of plant roots following the decapitation of the stem cell niche treated with BL (A), BL+IAA (B) or kyn and BRZ (C). n=50,40,95 for each condition in (A), (B) and (C), respectively. P values are according to Fisher exact test.

### Ectopic expression of *bin2-1* as a tool for tissue-specific inhibition of BR signaling

After establishing that the dual effect of BR is essential for meristem function and that the epidermis is an important tissue for BR-mediated auxin levels and response, we next asked if genes perturbed by BR and by auxin are commonly enriched in this tissue by 12h. DEGs were commonly noted in the outer tissues, i.e., epidermis and lateral root cap and columella (LR/C, Figure 2G). To assess whether BR signaling in the outer tissues is essential for meristem maintenance, we established a collection of wild type lines with targeted *bin2-1* expression to selectively inhibit the BR pathway in specific tissues. This included lines with *bin2-1* fused to the fluorescent markers Ypet or NeonGreen expressed in the epidermis (*pWER*, see also supplementary methods), LR/C (*pBRN2*), endodermis and QC (*pSCR*), stele (*pSHR*), and QC and initials (*pWOX5*) (Figure 5A). Remarkable, *bin2-1* expression in the outer tissues (*pWER* and *pBRN2*) featured a *bri1*-like meristem, manifested by wide meristems as in seedlings grown in the presence of BRZ (Figure 5G,H). Moreover, their width was unaffected by the addition of BL (Figure 5G,H, Figure S7). In addition, *pWER-bin2-1* lines had reduced root waviness in response to BL (Figure 5I). Intriguingly, stele *bin2-1* roots had the opposite phenotype, with a thinner meristem bearing longer meristematic cells, reminiscent of a high BR effect (Figure 5A,-C, Figure S7). BR biosynthesis genes have diverse expression patterns in the root, including expression in stele cells (Figure 5D, [11]) although these patterns are not necessarily overlapping in cell types composing it (http://bar.utoronto.ca/ based on [39]). As an example, *pCPD-NG-CPDter* established here demonstrates a specific predominant BR expression pattern in stele cells (Figure 5E). In addition, the transcription of BR biosynthesis genes are locally regulated by BRI1 in their corresponding tissues, where they are elevated in its absence [11]. Thus, it was hypothesized that specifically blocking BR signaling in the stele, as in *pSHR-bin2-1*, will alleviate an otherwise local transcriptional BR-driven repression of BR biosynthesis genes.

**Figure 5.**
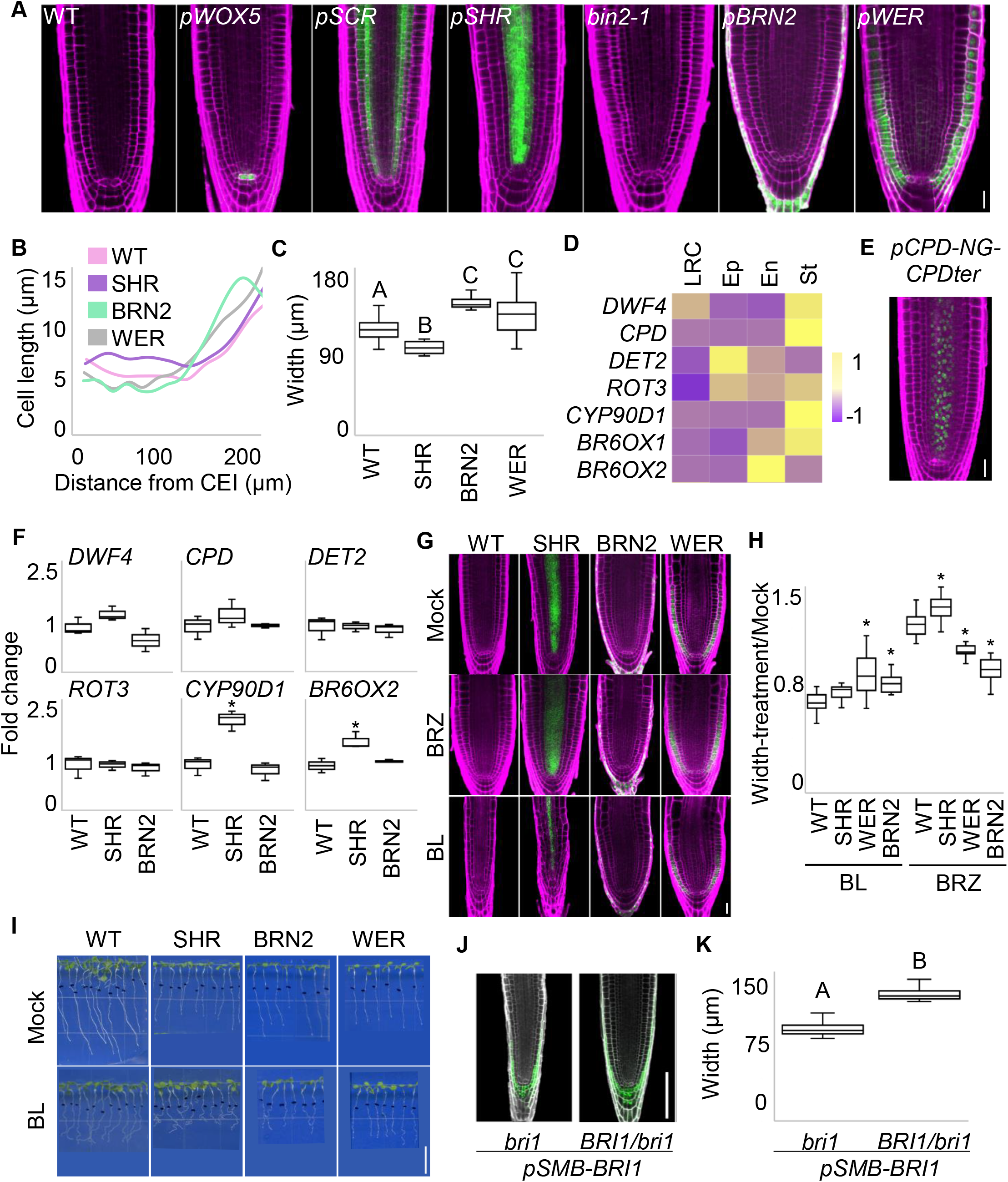
Tissue-specific *bin2-1* expression demonstrates that stele-BR controls its own homeostasis and modulates the meristem. (A) Confocal microscopy images of representative root meristems of 7-day-old wild-type, *BIN2/bin2-1*, and transgenic lines expressing *bin2-1-GFP* or *bin2-1-NG* (green) in different tissues driven by the following tissue specific promoters; *pWER, pBRN2, pSCR, pSHR* and *pWOX5*, PI signal: magenta (see methods for additional information). Note the opposing meristem phenotype of *pWER* and *pBRN2* and *pSHR*. (B) Average cell length as distance from CEI (cortex endodermis initial), in wild type and select *bin2-1* lines (See also Figure S7) (n≥8). (C) Meristem width as measured in lines as in (B). (D) Heatmap presenting relative expression levels of BR biosynthesis genes in different root tissues. Data based on [11]. (E) Confocal image of *pCPD-NG-pCPDter* meristem, CPD, green; PI, magenta. 7-day-old seedling is shown. (F) Relative levels of BR biosynthesis gene transcripts, as determined by real-time PCR analysis of RNA from meristems of 6-day-old roots of wild type and select *bin2-1* lines. Expression was normalized to that of wild type under mock conditions (n = 3 biologically independent samples for each line and treatment). (G) Confocal microscopy image of wild type and *bin2-1* lines *pWER*, *pBRN2*, and *pSHR*. Seedlings were germinated on mock or 3 μM BRZ, for 3 days, and then transferred to treatment plates of mock, 0.8 nM BL or 3 μM BRZ. Note the meristem insensitivity of *pWER* and *pBRN2* and the hypersensitivity of *pSHR* to BL. (H,I) Meristem width presented as ratio between treatment (BL or BRZ) to mock (n=65,65,55 and 43 for wild type, SHR, BRN2 and WER respectively) (H) and seedling phenotype (I) of lines as in (G). (J,K) Confocal microscopy image of the meristem of *pSMB-BRI1-Ypet* in the *bri1* mutant background and in the segregating *BRI1/bri1* background, and their corresponding width (n=20). Note the BL-like phenotype of *pSMB-BRI1* only in the *bri1* background. Asterisk or different letters indicate values with statistically significant differences (p ≤ 0.05, according to Tukey HSD). In A, E and G, Scale bars = 25μm. In I, Scale bar = 1cm.

### Inhibition of BR signaling in the stele elevates BR biosynthesis genes and sensitizes the root meristem to BR

To test the hypothesis that *pSHR-bin2-1* has higher transcript levels of BR biosynthesis genes we performed a real-time PCR analysis. The basal expression level of BR biosynthesis genes was 1.2-to 2-fold higher in root tips of *pSHR-bin2-1*, with significantly higher levels of *CYP90D1* and *BR6OX2/CYP85A2* transcripts than wild type (Figures 5F). In *pBRN2-bin2-1*, select genes had somewhat lower expression levels as compared to wild type (Figures 5F). The overall gene modulation in *pSHR-bin2-1* may lead to higher endogenous BR levels, perceived by BRI1 outside the stele. If this is the case, inhibition of BR production should suppress the high BR-like phenotype of *pSHR-bin2-1*, while addition of BL should enhance it. Indeed, meristems of *pSHR-bin2-1* seedlings grown in the presence of BRZ had a BR-deficient-like phenotype as wild type grown in BRZ (Figure 5G,H). Moreover, these lines were hypersensitive to BL, as suggested by their narrow and deformed meristems and enhanced waviness of roots as compared to wild type (Figures 5G,H,I). This hypersensitivity is likely sensed by the outer tissues. *pSMB-BRI1* in the *bri1* mutant background line was produced to direct BRI1 expression to the outer LR/C tissue (Figure 5J). Intriguingly, the plants had a narrow meristem (i.e., in accordance with high BR signaling) in the *bri1* mutant background but not in the segregating *BRI1/bri1* background (Figure 5J,K). In agreement with the hypersensitivity response of *pSHR-bin2-1* lines sensed in the outer tissues, *YUC9* elevation by BL exceeded that of wild type (Figure 6A). In conclusion, this tissue-specific *“bin2-1* tool” underscores BR homeostasis as an important determinant of meristem morphology and suggests that this control is decoded by outer tissues.

**Figure 6.**
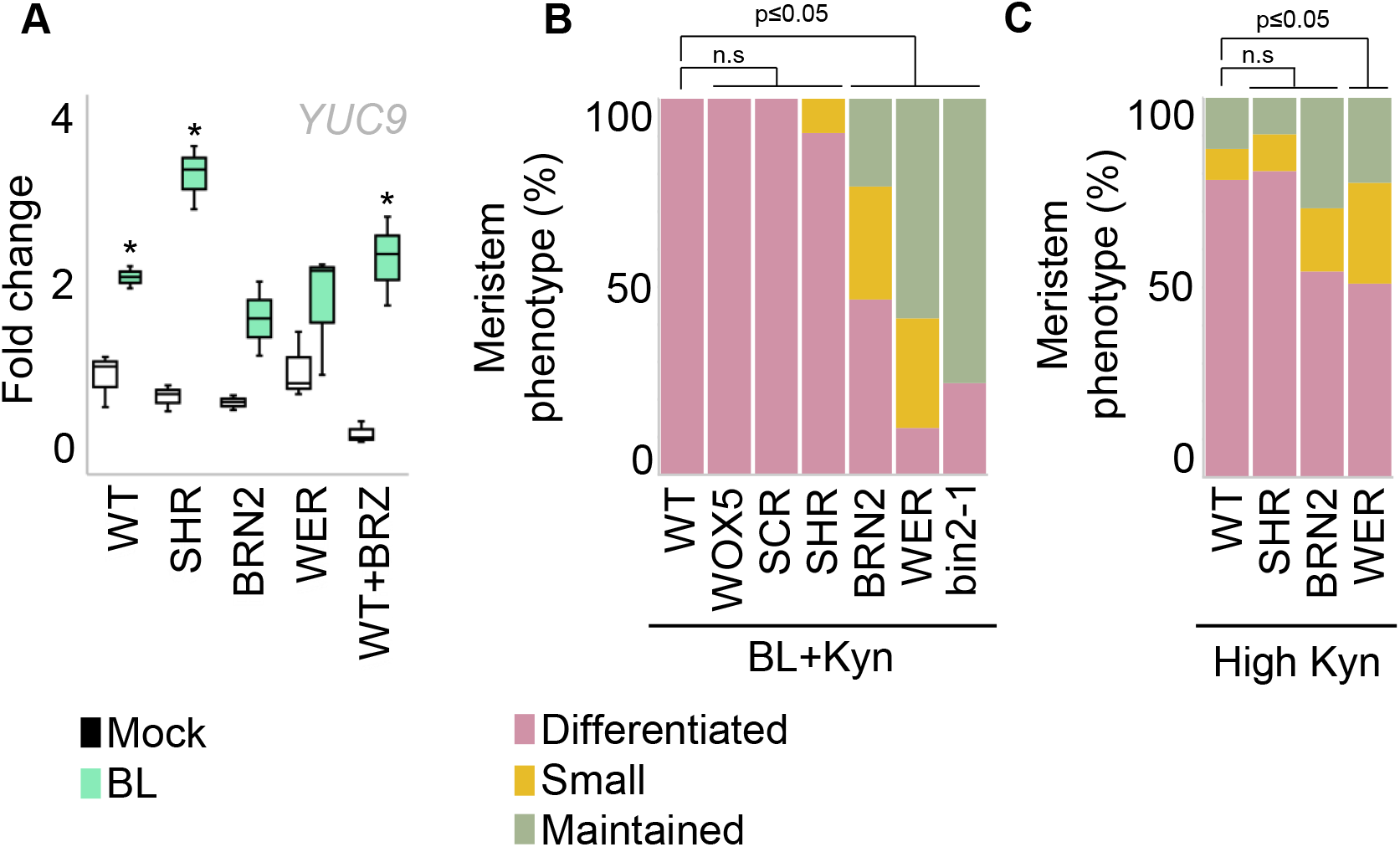
Inhibition of BR signaling in the outer tissues renders meristem insensitivity to both BR and low auxin. (A) Distribution of meristematic phenotypes in response to exposure to a combination of BL (0.8 nM) and Kyn (4 μM) for 96h (See also Figure S5) (n=58,37,17,18,22,24 and 30 for wild type, *bin2-1*, WOX5, SCR, SHR, BRN2 and WER respectively). (B) Relative *YUC9* transcript levels as determined by real-time PCR analysis of RNA from meristems of wild type and select tissue-specific *bin2-1* lines, following exposure to BL (0.8 nM) for 12h. Expression was normalized to that of wild type with mock treatment (n = 3 biologically independent samples for each line and each treatment). (C) Distribution of meristematic phenotypes of 9-day-old seedlings, in response to exposure to high 50 μM Kyn for 6 days (n=37,49,24 and 31 for wild type, WER, BRN2 and SHR respectively). In A and C, p values are according to Fisher exact test. In B, Asterisk indicates values with statistically significant differences between mock and BL (p ≤ 0.05, according to Tukey HSD).

### Inhibition of BR signaling in the outer tissues renders meristem insensitivity to both BR and low auxin

*bin2-1* expression in the outer tissues blocked *YUC9* elevation in response to BL (Figure 6A), suggesting that the outer tissues are necessary for the dual effect of BR on auxin. To determine whether blockage of BR signaling in the outer tissues also affects meristem response to auxin, *bin2-1* transgenic lines were subjected to the combined BL and Kyn treatment. Blocking BR signaling in the epidermis, and to some extent in the LR/C largely maintained an intact meristem (Figure 6B, Figure S7), while all other *bin2-1* lines tested (i.e., QC and initials, endodermis and stele) showed meristem differentiation phenotypes. Furthermore, inhibition of BR signaling in the epidermis and LR/C was less affected by low auxin compared to its inhibition in the stele and as compared to the wild type (Figure 6C). Thus, BR signaling in the outer tissues is necessary for its dual effect on auxin, and likely depends on BR homeostasis in the inner tissues.

## Discussion

### The root meristem is sustained by a dual BR effect on auxin

The significance of complex intertwining of signaling pathways lying at the heart of biological systems is often inherently challenging to decipher. By quantifying and interfering with an incoherent effect of BR on auxin, we uncovered its essential role for maintaining a functional root meristem (Figure 7). BR elevated the YUC9 reporter line in the epidermis, corroborating earlier findings that showed epidermal elevation of this gene and other auxin biosynthesis genes [11]. Moreover, we established here that BR elevates auxin levels in the root tip. In agreement, inhibition of auxin synthesis was largely epistatic to the effect of BR on the elevation of nuclear auxin signaling input. This also aligned with the observation that epidermal BRI1 overexpression was sufficient to elevate auxin maxima zone in the meristem [11]. Together, BR elevates auxin signaling input by controlling YUC gene(s) expression and auxin levels and by controlling the levels of PILs [12]. In parallel to BR mediation of auxin synthesis and consequently of auxin signaling input, the transcriptional readout for auxin output is repressed in the epidermis and the inner tissues. Overall, this BR-mediated repression of auxin transcriptional output is in agreement with a previously reported whole-root RNA-seq analysis which demonstrated that exogenous application of IAA and BL have opposing transcriptional effects on most of the DEGs [14].

**Figure 7.**
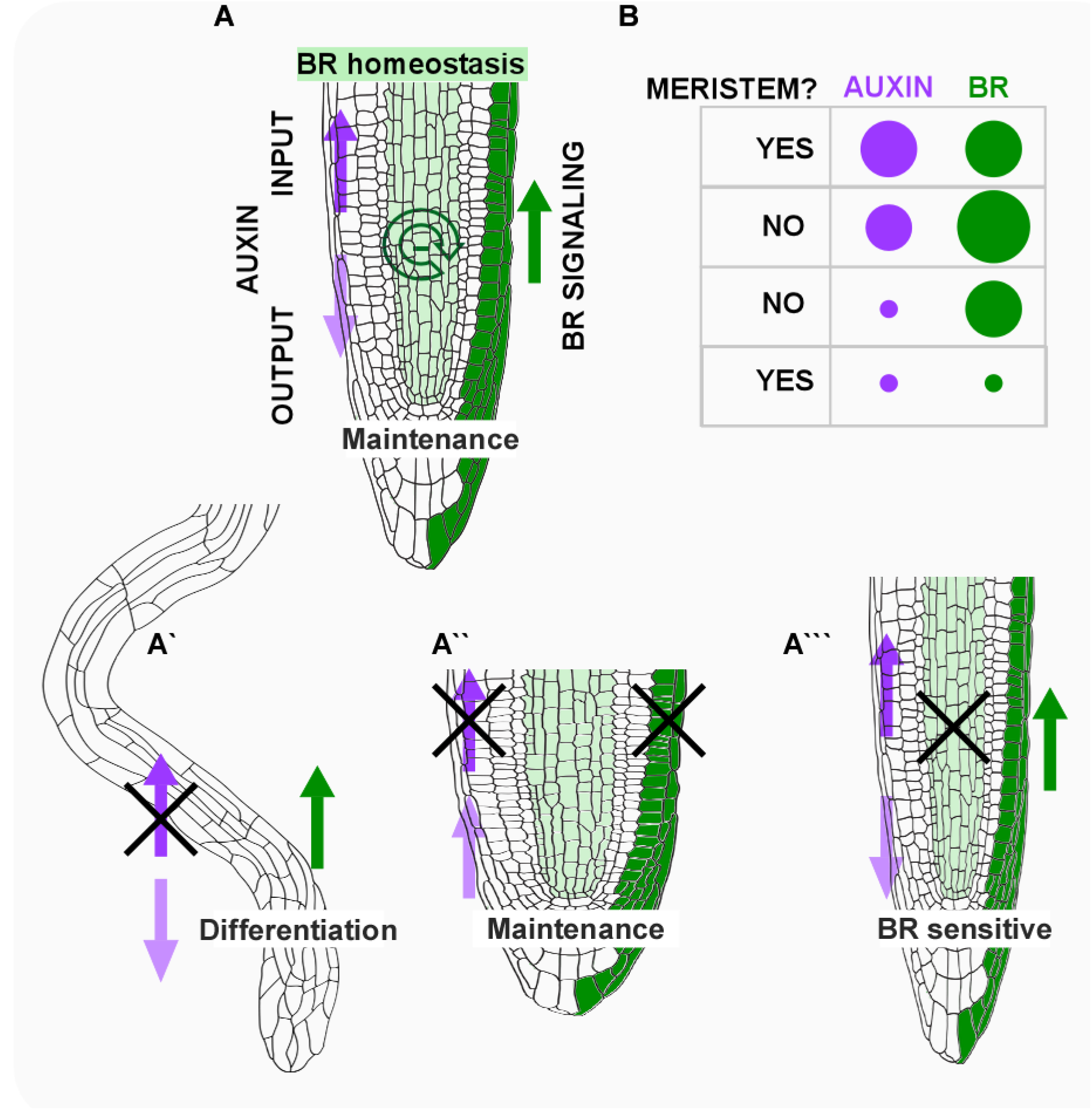
BR dictates auxin requirement for meristematic state: a proposed mechanism. (A) The dual effect of BR signaling, that both promotes auxin synthesis and opposes auxin signaling output is critical for meristem maintenance. BR signaling in the outer tissues is essential for this duality while BR activity in inner tissues is necessary to control BR homeostasis. If BR-mediated elevation of auxin synthesis is blocked, the meristem undergoes fast differentiation (A’); low BR rescues meristem loss caused by a severe reduction in auxin synthesis, accompanied by elevation of auxin output (A”); blocking BR signaling in the stele relieves the negative feedback on BR biosynthesis genes, sensitizing the meristem to BR. The outcome of these perturbations points to a critical ratio between BR and auxin (and not an absolute level, represented as circle size) that supports a meristematic state (B).

The present study suggested that the resistance of the meristem to hormonal fluctuation relies on BR-mediated elevation of auxin. If only one hormone is perturbed, i.e., if auxin biosynthesis is moderately inhibited or BR is moderately elevated, meristem size is reduced but remains intact. However, a similar but concomitant perturbation of these hormones leads to rapid meristem loss through a differentiation process that becomes irreversible by 48h. Perturbing the ability of BR to elevate auxin, renders it an executer of the differentiation program. Indeed, it was accompanied by a massive impact on gene expression, among which only a small fraction of genes returned to basal levels, unlike the effect of a single hormone perturbation, suggesting a plausible loss of regulatory feedback. The resistance of the meristem to hormonal fluctuation shown here may involve additional layers of molecular and physiological interactions between BR and auxin which are thought to maintain a synchronized transcriptional response [12]. In the canonical auxin signaling pathway, the hormone binds to the transport inhibitor response 1/Auxin Signaling F-Box (TIR1/AFB) family of F-box receptors and triggers the degradation of Auxin/Indole-3-Acetic Acid (Aux/IAA) co-repressors. Consequently, Auxin Response Factors (ARFs) are activated and regulate gene expression [40]. Mechanistically, select ARFs and BES1/BZR1 can undergo similar post-translational modifications by BIN2 and by the SUMO group by proteases OTS1/OTS2 [41, 42], with opposing consequences on their transcriptional activities [43, 44]. Thus, these enzymes may regulate the differential transcription that preserves a meristematic state.

Regardless of the precise mechanism involved, maintenance of an intact meristem and capacity to regenerate upon deleterious low auxin levels, when BR was absent was quite striking. The presence of the auxin transcriptional output in the meristem could simply reflect enhanced sensitivity to auxin. Alternatively, a core transcriptional output could be still functional in the absence of both hormones. While it is uncertain if a generic program underlies meristem function, clearly an equivalent interplay between BR and auxin also dictates meristem regeneration after wounding. Injured root meristems rely solely on local auxin synthesis, as evidenced by loss of regeneration when auxin production is inhibited. However, our study demonstrated that loss of regeneration in these same conditions is largely recovered, if BR levels are also reduced. In agreement, addition of very low amount of BL blocked regeneration, an effect which was counteracted by addition of auxin. Overall, this work demonstrated, both in the context of an established meristem or of the *de novo* formed meristem, that a critical ratio between BR and auxin, irrespective of their levels, dictates the existence of the meristem (Figure 7).

### Inter-tissue communication controlling the meristem: negative feedback on BR production in the stele is linked to BR-decoding in the outer tissues

Tissue-specific BR decoding was primarily inferred from complementation experiments where BRI1 was targeted to specific cells and tissues, thereby limiting BR perception to tissues of interest [10, 45]. This approach revealed opposing BR signals in the outer and the inner tissues and their balancing effect on the meristem [10, 11, 13]. Here, we also provide mechanistic insight into this balance. Using a complementary tool which selectively inhibited BR signaling via expression of *bin2-1*, the present work uncovered an inter-tissue interplay of BR signaling, BR biosynthesis and BR homeostasis in the root meristem. BR signaling in the epidermis and lateral root cap was shown to be essential for regulation of meristem development, for the dual effect on auxin and for meristem sustainability. Specifically, lines expressing *bin2-1* in outer tissues had a *bri1*-like meristem phenotype and limited elevation of YUC9 upon exposure to the hormone and more roots maintained an intact meristem in response to low auxin. Importantly, this tool separated the response of the outer tissues from that of the stele, which exhibited an opposite meristem phenotype, with roots resembling those treated with moderate BR levels, and hypersensitive to the hormone (Figure 7). This was accompanied by elevation of several BR biosynthesis gene transcript. Some genes, e.g., DET2, DWF4/CYP90B1 and CPD/ CYP90A1, have non-redundant homologues in Arabidopsis and their loss-of-function result in a typical BR-deficient phenotype [46–48]. Others, like CYP90D1 and ROT3/ CYP90C1 and BR6OX1/ CYP85A1 and BR6OX2/ CYP85A2 (that involved in the last step of BR synthesis), have partially redundant functions [49, 50]. We confirmed, via a reporter line, that the non-redundant CPD gene is restricted to the stele tissue. As BRI1 was shown here to be required for gene expression in the outer tissues and the control of meristem size, BR precursors and active BR likely move this short distance. BR homeostasis in the meristem may also be regulated by catabolic genes [51] expressed in the columella and lateral root cap, as suggested by the elevated BZR1 nuclear levels in the QC cells in their absence [14]. The negative feedback of BR signaling on its own production is thought to be important, but perturbations that directly link it to development are inherently more difficult to demonstrate. Hence, an additional important aspect of this study is the realization that a spatiotemporal negative feedback is an important developmental determinant of meristem shape and maintenance.

## Methods

### Plant material, growth conditions and chemical treatments

All Arabidopsis (Arabidopsis thaliana) lines were in the Columbia-0 (Col-0) background. The following lines were used: *wei2,7* [37], *yuc3yuc5yuc8yuc9 (yuc-q)* [3], *cpd* [47], *bin2-1* [52], *bri1* (GABI_134E10) [53, 54]*, bri1-116, brl1,brl3 and bri1,bil1,bil3* (triple mutant) are described in [11], *DII-VENUS* [32], *DR5v2-ntdTomato–DR5-n3GFP* [35], *pIAAmotif-mScarlet-NLS*, modified from [36] and *pYUC9-VENUS* [38]. Plates with sterilized seeds were stratified at 4°C for 2-5 days in dark. Seedlings germinated on one-half-strength (0.5) Murashige and Skoog (MS) medium supplemented with 0.2% (w/v) sucrose, with the exception of root cutting-assay where 0.8% sucrose was used. Seedlings were transferred after 3 days to hormonal and chemical treatments and analyzed after additional 4 days (unless indicated otherwise). For short term treatments (as in Figure 1, Figure S1), seedlings were transferred for additional 12h. For long term treatments seedlings were transferred for additional 10-12 days (Figure 3D,E, Figure S5). For BRZ treatments, seeds were germinated on BRZ unless indicated otherwise (as in Figure 3D, Figure 5, Figure S5). All hormones and chemicals were dissolved in DMSO, that was used as mock treatment. Hormonal concentrations used were primarily 0.8nM BL (Chemiclones), 4μM Kyn (Sigma-Aldrich) and their combination unless indicated otherwise, 3μM BRZ [34], 0.2μM NAA (Sigma-Aldrich) and IAA (concentrations as indicated in the text, Sigma-Aldrich). Seedlings were grown in 16h light (approximately 70 μmol m^−2^ s^−1^)/ 8h dark cycles or continuous light, at 22°C.

### Constructs and transgenic lines

For tissue specific expression of BIN2-1, *bin2-1-GFP* was cloned as a KpnI/KpnI fragment into pBJ36-containing *pSCR, pSHR* as in [8] and *pBRN2* as in [11] and *pWOX5*. Promoter WOX5 was designed using 4985bp fragment upstream to the first ATG of the *WOX5* gene. The final constructs were subcloned into the *pMLBART* binary vector as NotI/NotI fragments. *pWER-BIN2-1-NeonGreen, pSMB-BRI1-Ypet* and *pCPD-N7-NeonGreen CPDterm* were generated using the Golden Gate MoClo Plant Tool Kit [55] (see supplementary methods). Final constructs were in *pICSL4723* harboring kanamycin selection. For cloning of *pGL2-YUC5, YUC5* was amplified from Col-0 genomic DNA. Amplified fragment was ligated into BJ36 vector containing *pGL2* sequence [8] using KpnI-BamHI restriction sites and GFP was then inserted using BamHI-BamHI sites. The final construct was subcloned into the *pMLBART* binary vector as NotI/NotI fragments. Plant transformation to wild-type Col-0, *BRI1/bri1* and *yuc-q* background was performed using *Agrobacterium tumefaciens* (GV3103)-mediated floral dip transformation method. Transgenic lines were selected for basta or kanamycin resistance (see notes in supplementary methods).

### Quantification of confocal images

For cumulative cell lengths, the length of each cortical cell along its corresponding cell file in the meristem was measured, and averaged in number of roots as indicated in each figure legend. Meristematic cell number was calculated according to average cumulative cell lengths along cortex cell files, determining meristem size as the average cell size that doubled its size from the initial cortical cell measured. Meristem width was measured as width of the root meristem at 150 μm from the QC. The fluorescence signal intensity (mean gray value) of the presented markers was quantified using the Fiji software [56]. The relevant region quantified is indicated in each figure legend. For quantification of the epidermal fluorescence signal intensity along the meristem, confocal images of the epidermal plane were analyzed, measuring signal intensity (mean gray value) along the epidermal cell file, signals lower than 10 were discarded from the analysis. Log10 values of the signals were smoothed (loess, [57]) by their distance from the tip and plotted in a graph. In Figure S1, similar analysis was performed but the threshold to discard background signal intensity was 3100 (mean grey value).

### Root-cutting assay

Seven-day-old seedlings were cut at 120 μm from the tip of the meristem as described in [58] and moved to recovery plates, with or without treatments. The frequency of regeneration was evaluated 3 days after cutting. Between three to five independent cutting sessions (batches) were used for each experiment, each with a matching wild type or mock control.

### IAA measurements

For IAA measurement as in Figure 1C, root tips of 6-day-old seedlings, after 12h of 0.8 nM BL, were harvested and immediately frozen in liquid nitrogen until further extraction, in five biological repetitions. For IAA measurement as in Figure 3C, whole roots of 7-day-old seedlings after hormonal or chemical treatment were harvested and immediately frozen in liquid nitrogen until further extraction, in three biological repetitions.

IAA was measured by UPLC-ESI-MS/MS, as described in supplementary methods.

### RNA extraction, real-time PCR and sequencing and data analysis

Total RNA was extracted from roots (cut at about 0.2-0.5 cm from the tip) using the Spectrum plant total RNA kit (Sigma). RNA sequencing and analysis was performed at the Technion Genome Center (LSE). Samples (3 biological replicates) were prepared using CEL-Seq2 sample preparation protocol [59]. Samples were sequenced on Illumina HiSeq 2500. Reads were aligned to the TAIR10 assembly of the Arabidopsis thaliana genome, using Tophat2 version 2.1.0. Transcript coordinates from the TAIR10 reference set were used to guide the alignment process. After alignment, the numbers of reads per gene were estimated using HTseq-count version 0.6.1. DESeq2 R package version 1.18.1 was used to normalize read counts and perform pair-wise differential expression analysis. Gene sets were identified based on their differential expression. Significantly modulated genes were considered with a fold change of 1.5 and adjusted p-value (false discovery rate) of 0.05 or less, comparing the hormonal perturbations to the control in the same treatment duration (12, 24 or 48h). Quantitative real-time PCR assays were performed as in [8]. Experiment conducted in three biologically independent repeats.

### Confocal Microscopy

Confocal microscopy was performed using a Zeiss LSM 510 (Zeiss, Jena, Germany) confocal laser scanning microscope with a LD LCI Plan-Apochromat 25× water immersion objective (NA-0.8). Roots were imaged in water, or with water supplemented with Propidium iodide (PI, 10 μg/mL). The green fluorescence protein (GFP, NeonGreen) and PI were excited by Argon laser (488 nm) and by DPSS laser (561 nm), respectively. Fluorescence emission signals for GFP/NeonGreen and for PI were collected by PMT detectors, with band pass filter (500-530 nm) and with long pass filter (575 nm), respectively. Venus fluorophore was excited by Argon laser (488) nm and the emission was detected with band pass filter (500-530nm). mScarlet fluorophore was excited by DPSS laser (561 nm) and the emission was detected with long pass filter (575 nm). All image data were analyzed using Fiji software.

### Statistical analysis

Data was analyzed by one-way analysis of variance (ANOVA) using JMP Pro version 14.0.0 software (SAS Institute, Cary, NC, USA). Differences between treatments were determined by Tukey–Kramer HSD test (p≤0.05), for multiple comparisons. Differences between meristem phenotype distribution (as in Figure 3,4 and 6) was determined by Fisher exact test, for 2X3 tables the Freeman-Halton extension [60] was applied, as detailed in the figure legends.

## Acknowledgments

We thank Y. Zhao, J. Li and E. Russinova for sharing seed material, T. Asami for providing BRZ, F. Abd el Azez, A. Buri, Y. Ben Eliahu, L. Nauman, A. Farhat, D. Eisler, N. Holland and E. Lavert for their excellent technical assistance, the Life Sciences and Engineering Infrastructure Center (N. Dahan, Y. Lupu-Haber and the Genome Center), the Russell Barrie Nanotechnology Institute at the Technion and the Metabolic Profiling Unit at the Weizmann Institute (S. Malitsky, I. Panizel). We also thank Y. Eshed for helpful comments on this study. This research was supported by the Israel Science Foundation (grant No. 1725/18).

