## Supplementary Material for "Auxin requirements for a meristematic state in roots depend on a dual brassinosteroid function"

**Supplemental Information**

**Supplemental Figures**

Figure S1

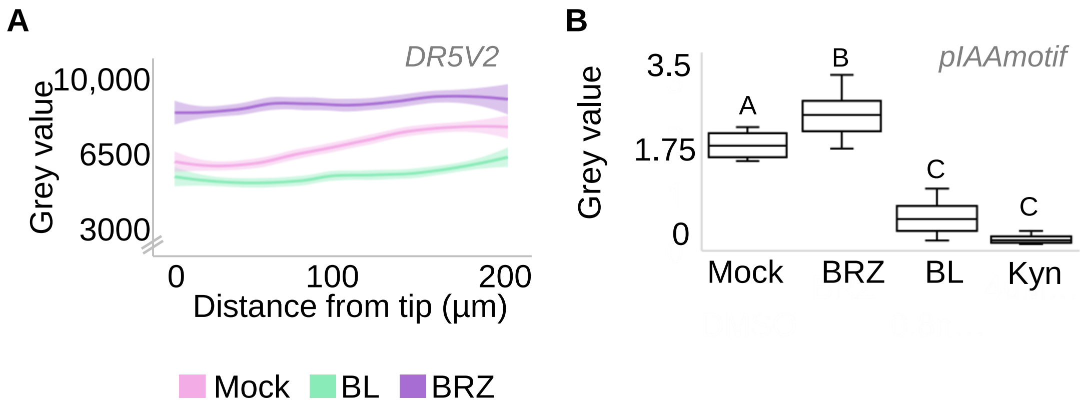

**Figure S1. BR inhibits auxin signaling output**

(A) Quantification of the fluorescence signal in the epidermis along the meristem of DR5v2-ntdTomato roots treated with mock, 0.8 nM BL or 3 µM BRZ, for 12 h (n=15,12,22 respectively).

(B) Quantification of the fluorescence signal ($log_{10}$ scale) in the stele of *pIAAmotif-mScarlet-NLS* roots treated with mock, 0. 8nM BL or 3 µM BRZ and 4µM Kyn for 12h (n=15,12,14,11 respectively), as in Figure 1.

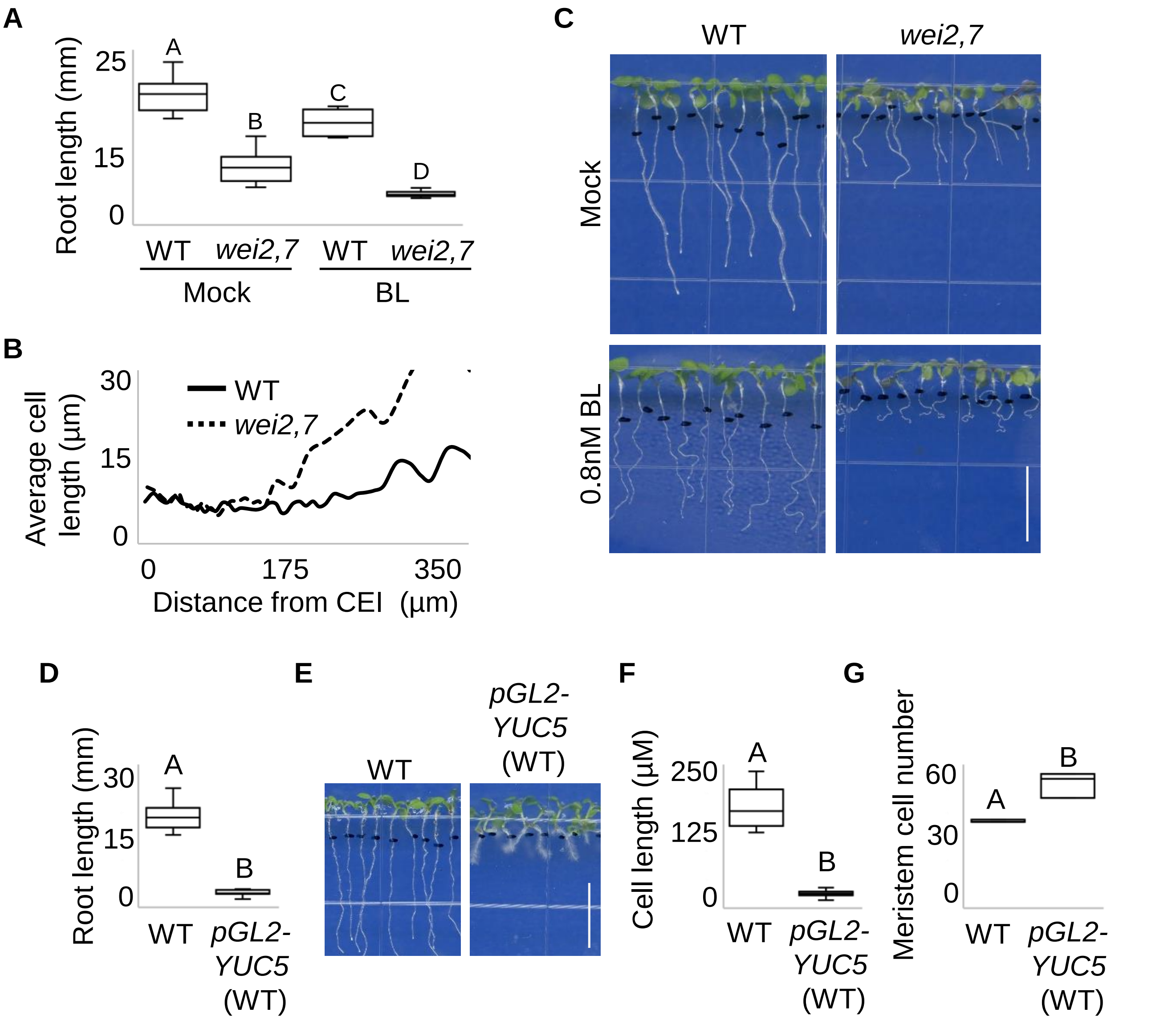

**Figure S2. The root meristem of auxin biosynthesis mutant is hypersensitive to BR while epidermal expression of YUC is sufficient to confer high auxin-like phenotype**

(A-C) Phenotype of wild-type and *wei2,7* seedlings after transfer to mock or BL (0.8 nM), for 96h and phenotype of wild type and GL2-YUC5 lines (D-G). (A) Root length (n=15) (B) average cell length versus position from cortex endodermis initial (CEI) (n=8), and (C) whole-seedling phenotype. (D) Root length (n=10) (E) whole-seedling phenotype, (F) meristematic cell length (n=16 and 37 for WT and GL2-YUC5 respectively) and (G) meristem cell number (n=5). The different letters indicate values with statistically significant differences ($p \leq 0.05$). Scale bar = 1cm.

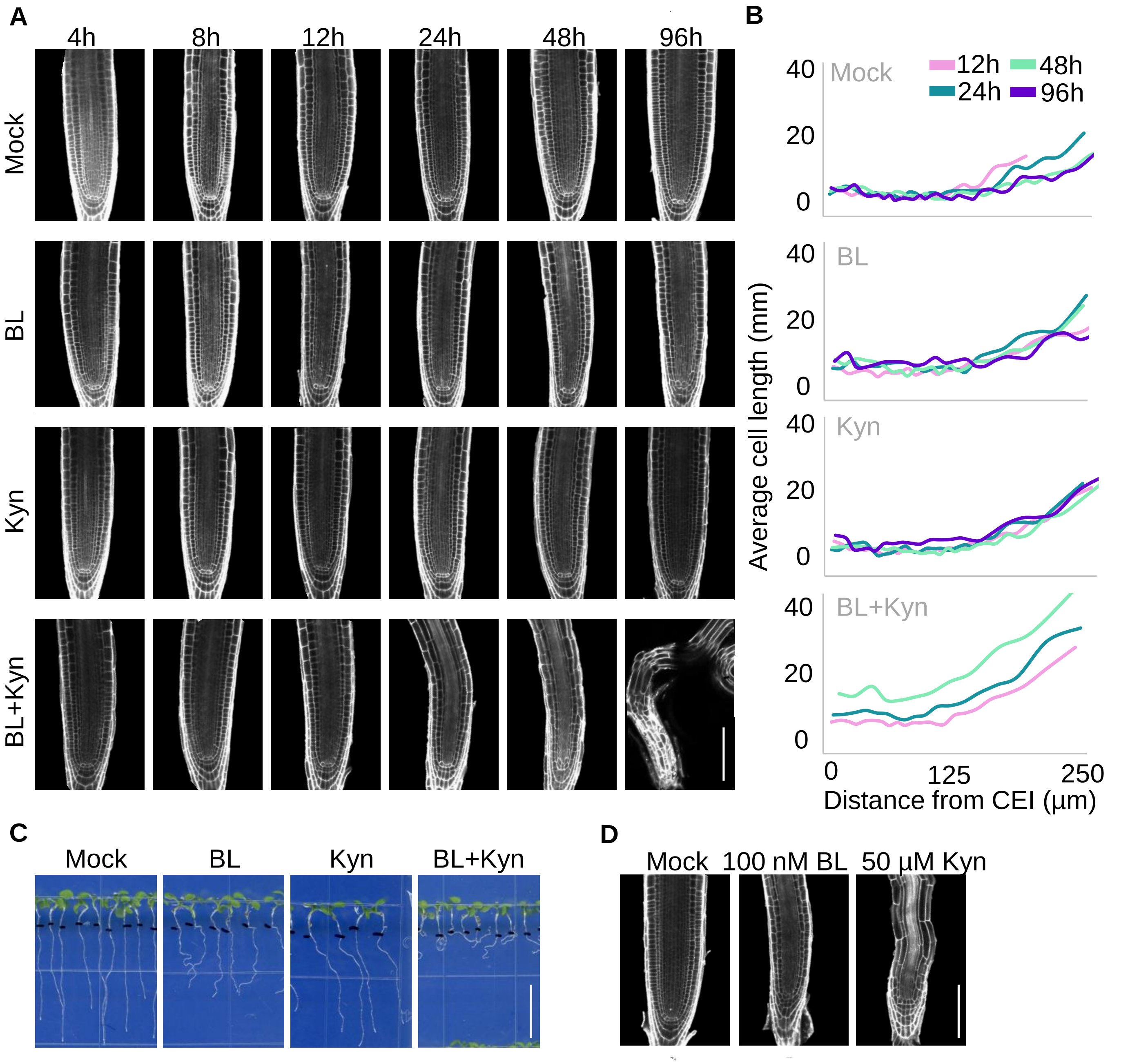

**Figure S3. Dynamics of meristem differentiation in response to hormonal perturbations**

(A) Confocal microscopy images of representative root meristems at 4h-96h after transfer to mock, 0.8 nM BL, 4 µM Kyn or their combination.

(B) Average cell length versus position from CEI (cortex endodermis initial), of seedlings as in (A); see also Figure 2 (n≥8).

(C) Phenotype of 7-day-old seedlings grown as in (A), 96h after transfer to hormonal treatments.

(D) Confocal microscopy images of representative root meristems treated with 100 nM BL or 50 µMu Kyn. Scale bars =100 µm in A and D, scale bar = 1cm in C.

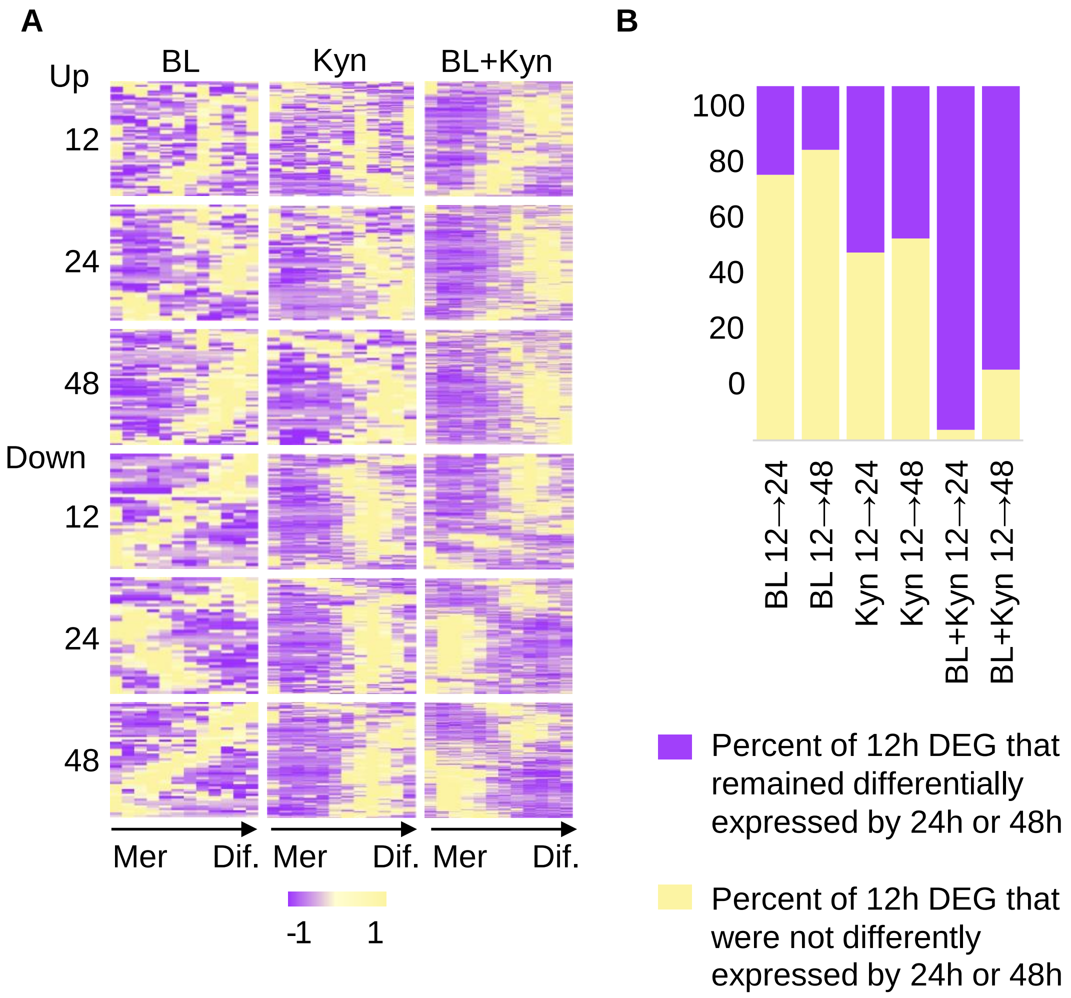

**Figure S4. Simultaneous hormonal perturbation is accompanied by deregulation of gene expression**

(A) DEGs in different root zones at 12 h, 24 h and 48 h. Note that combined BL and Kyn treatment caused concomitant loss of meristematic genes and elevation of differentiation genes by 12h.

(B) Percent of DEGs after 12 h treatment ($p \leq 0.05$, and FC≥ 1.5) that are not differentially expressed at 24 h or 48 h of treatment (see also Figure 2).

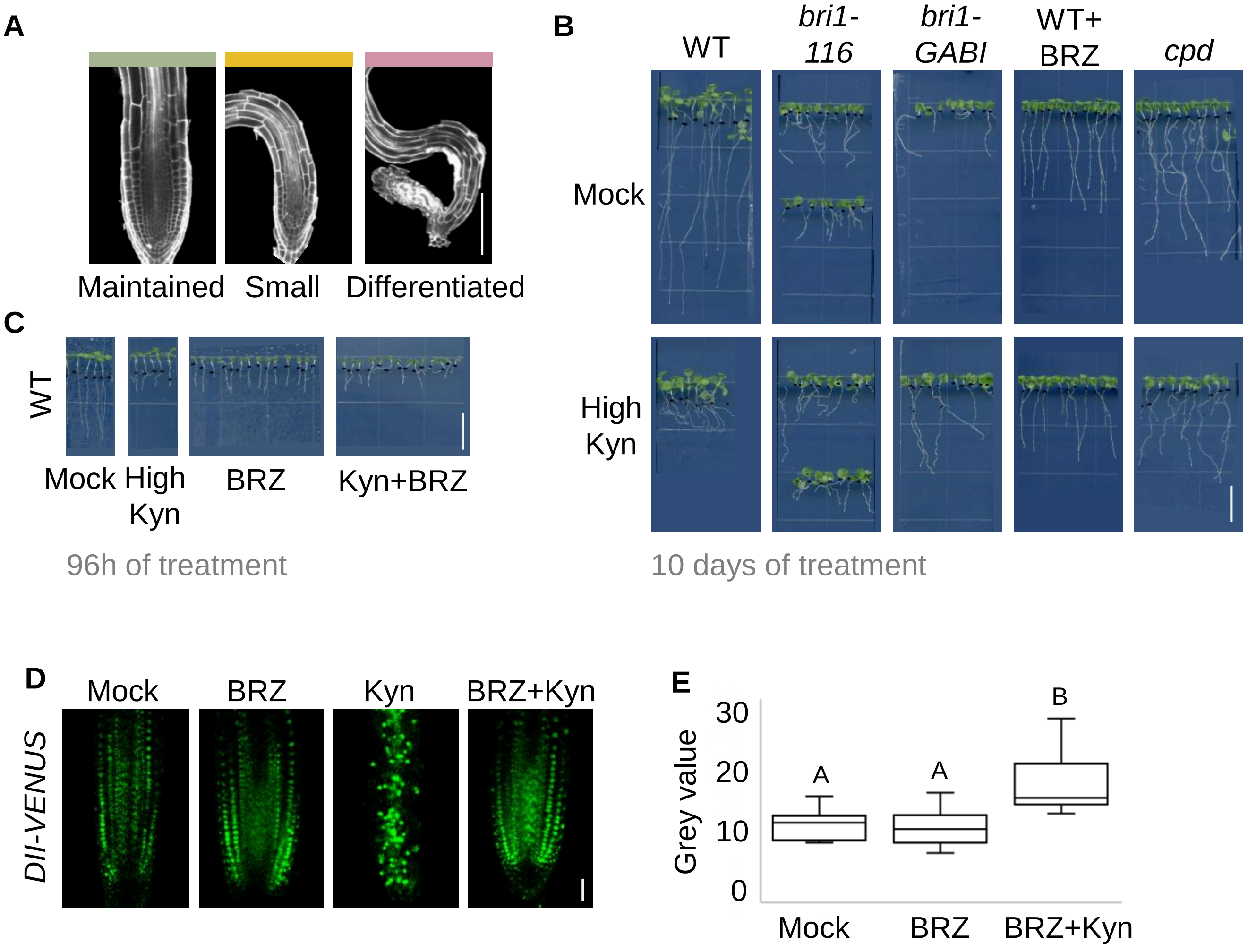

**Figure S5. The root meristem of BR-deficient mutants is maintained upon a severe block of auxin biosynthesis**

(A) Confocal microscopy images showing the range of meristem phenotype following high-Kyn or combined BL and Kyn treatment. (See Figure 3 and Figure 6)

(B) Seedlings of wild-type, *bri1-116, bri1-GABI* and *cpd,* 10 days after transfer to mock or high Kyn (50 µM). Wild type seedlings were also germinated on BRZ and then subjected to combined BRZ and high Kyn treatment.

(C) 7-day-old wild-type seedlings that were germinated in the presence or absence of 3 µM BRZ before being transferred to high Kyn conditions, for 96 h. In B and C, scale bar = 1cm.

(D,E) Confocal microscopy images and corresponding quantification of *DII-VENUS* 10 days after transfer to mock, 3 µM BRZ, high Kyn (50 µM) or high Kyn+BRZ (n=10), scale bar= 25 µm.

Figure S6

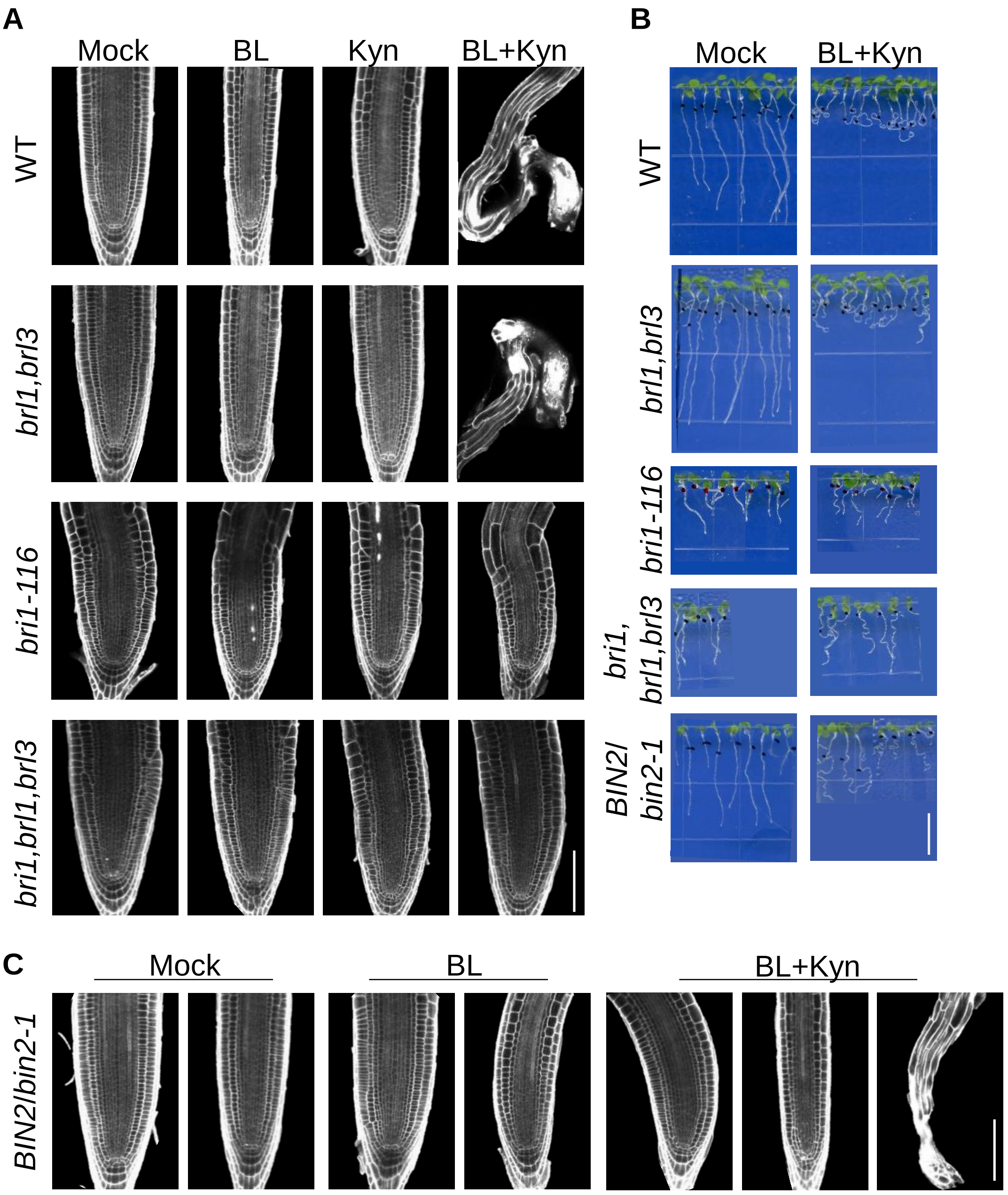

Meristem phenotype

| | | | | Total |
|---|---|---|---|---|
| WT | 48 | - | - | 48 |
| *BIN2/bin2-1* | 9 | - | 28 | 37 |

Differentiated
Small
Maintained

**Figure S6. The root meristem of BR-deficient mutants is maintained upon concomitant moderate perturbation of BR and auxin**

(A,B) Confocal images (A) and seedlings phenotype (B) of wild type, *bri1-116, brl1,brl3 and bri1,brl1,brl3,* 96 h after transfer to mock, BL, Kyn or their combination.

(C) Confocal images of *BIN2/bin2-1* subjected to hormonal treatments as in (A,B). The *BIN2/bin2-1* population segregated to roots, with meristem insensitivity to BL, as indicated by the wide meristem phenotype (Left root). *BIN2/bin2-1* meristem phenotypes following BL and Kyn treatment for 96h are summarized in a table, see also Figure 6. Scale bars =100 µm in A and C, scale bar = 1cm in B.

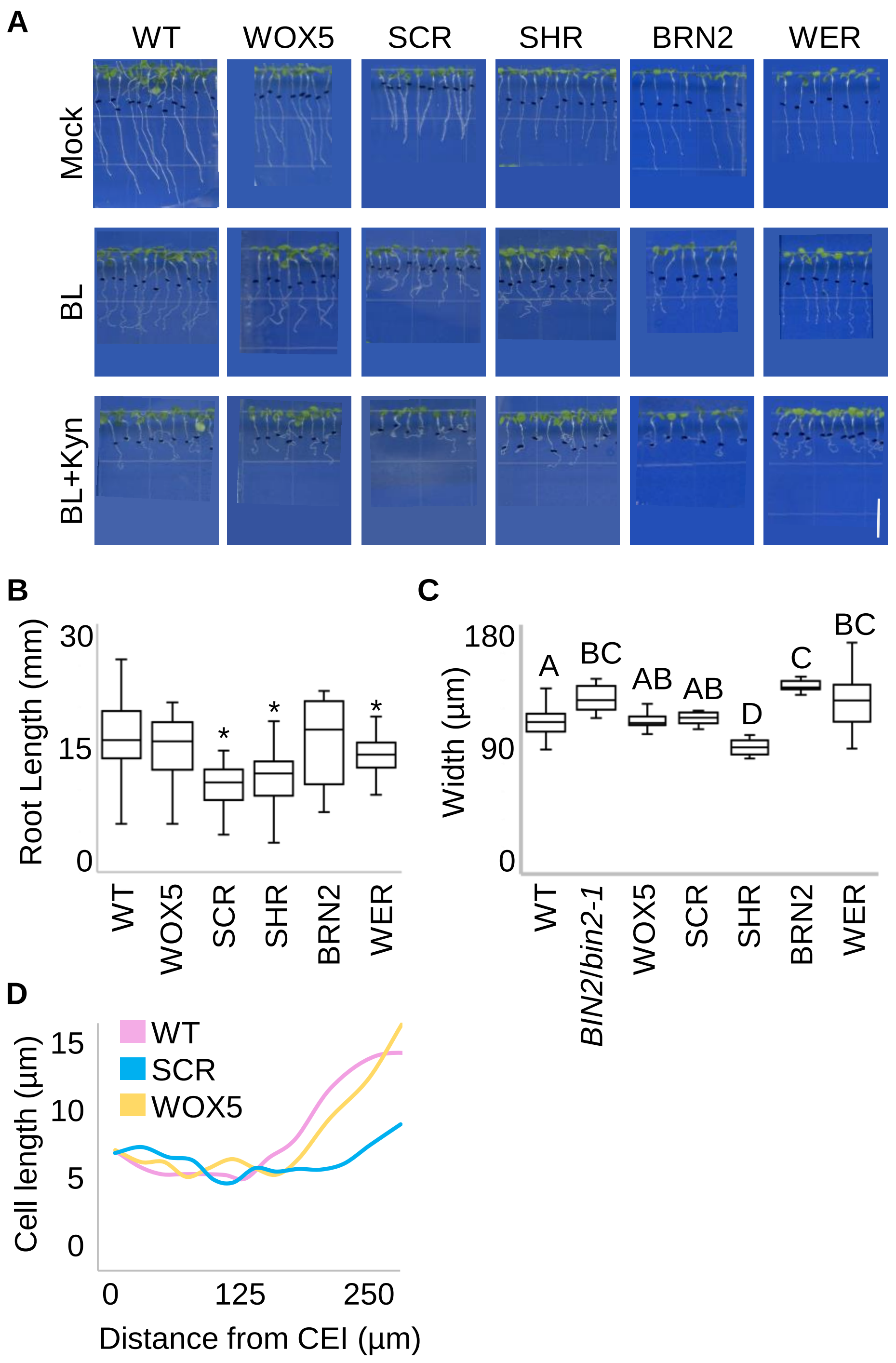

**Figure S7- Phenotypes of transgenic lines with tissue-specific *bin2-1* expression**

(A) Seedling phenotype of wild type and transgenic lines expressing *bin2-1* driven by select tissue-specific promoters: *pWER, pBRN2, pSCR, pSHR and pWOX5*, 96 h after transfer to mock, BL, kyn or their combination (experiment as in figure 5).

(B) Root length of seedlings as in (A) (n≥15).

(C-D) Meristem width (n≥15) (C) and meristematic cortical cell length, measured as distance from CEI (cortex endodermis initial) (D) (n≥8) of wild-type and transgenic lines expressing *bin2-1* in specific root tissues. The different letters indicate values with statistically significant differences between the different lines. The asterisk indicates values that were statistically significantly different from those of the wild type, both according to Tukey HSD ($p \leq 0.05$) (C and D are elaborated experiment as in Figure 5).

### Supplemental Experimental Procedures

#### Constructs and transgenic lines:

For *pWER-BIN2-1-NeonGreen, pWER* was designed using the 3730 bp fragment upstream to the first ATG. *pWER* in level 0 (in *pICH41295*) was then sub cloned along with additional level 0 parts: *bin2-1* (in *pAGM1287*), NeonGreen (in *pAGM1301*) and WER terminator (in *pICH41276*) into level 1 (*pICH47742*). The constructed level 1 was then subcloned to level 2 construct (*pICSL4723*), together with a level 1 kanamycin resistance gene (*pICH47732*). For *pSMB-BRI1-Ypet*, *pSMB* in level 0 [1] was sub cloned along with, *BRI1* (in *pAGM1287*), *Ypet* (in *pAGM1301*) and *rbcs* terminator (in *pICH41276*) into level 1 construct (*pICH47742*). The constructed level 1 was then subcloned with additional level 1-components as above, into level 2 construct (*pICSL4723*). For *pCPD:N7:NeonGreen:CPDterm*, *pCPD* was designed using 2Kb fragment upstream to the first ATG of the *CPD* gene .*pCPD* in level 0 (pICH41295) was sub cloned along with NeonGreen (in pAGM1301) and CPD terminator (in pICH41276) into level 1 construct (pICH47742). The constructed level 1 was then subcloned with additional level 1-components as above, in *pICSL4723* . Recognition sequences for the Golden Gate based type IIs enzymes were eliminated if they were present in cloned fragments. Primers are listed in table S2. For each construct used, 2-3 independent transgenic lines were generated, all showing similar phenotype. All *pSCR* lines showed strong expression both in the endodermis and cortex. The *pSCR* line with strongest expression was chosen for this study, also showing a smaller shoot compared to wild type. In the case of the *pWER* lines, all lines showed faint fluorescent signal also in the cortex. The *pWER* line used in this study showed stable epidermal fluorescence signal, but sporadically completely lost its signal, likely a result of gene silencing. In cloning of *GL2-YUC5*, since *yuc-q* is resistant to basta, transgenic *GL2-YUC5* in *yuc-q* were screened based on short and hairy root phenotype and then verified by PCR.

#### IAA measurements

Auxin extraction was carried out based on the method for phytohormones described in [2, 3] in the Metabolic Profiling Unit (Weizmann Institute). 150 mg of ground frozen plant tissue was extracted overnight at -20°C with 1 ml methanol/water/formic acid (15/4/1 v/v/v), containing 25 ul of 1 ug/ml d5-IAA (Sigma) as stable isotope labeled internal standard (IS), followed by second extraction with 0.5 ml of the same solvent without IS. After evaporation under nitrogen stream and reconstitution to 1 ml 1% acetic acid in ddw, the extract was applied on Oasis MCX 1ml/30cc cartridge preconditioned with 1 ml ACN, 1 ml MeOH, 0.5 ml HCl 0.1M and 1 ml 1% acetic acid. After loading the samples and washing the cartridges with 1 ml acetic acid 1%, auxins were eluted with 1.5 ml MeOH, evaporated to dryness under nitrogen stream and reconstituted in 50 ul 5% ACN in ddw. Auxins were measured by UPLC-ESI-MS/MS equipped with Waters Acquity UPLC system with quaternary pump and Acquity UPLC BEH C18 column (1.7 um,2.1x100mm, Waters). Auxins were separated at a flow rate of 0.3 ml/min with gradient of 0.1% acetic acid in 3% ACN in ddw (A) and in 100% ACN (B). At first minute of the injection, 5% B was run, followed by 1-minute gradient to 15% B, 10.5-min gradient to 45% B, then 0.5 min gradient to 100% B and 1.5 min isocratic wash with 100%B. After that, the system was returned to the initial condition (95% A) in 0.25 min, and equilibrated during 0.25 min. The autosampler was cooled to 10°C and the column heated to 35°C. MS detector (Waters TQ-XS) was equipped with ESI source used in positive mode, capillary voltage 2 kV. The measurement was performed in MRM mode, 2 MRM traces for each compound - one for quantification, and another for identification. MS parameters for the MRM transitions, together with standard names, abbreviations and source are summarized in Supplementary. Data were processed with MassLynx software with Targetlynx (Waters). Quantification of IAA was done against external calibration curve, using IAA/d5-IAA peak ratios. Other auxins were quantified relatively, using d5-IAA as internal standard.

**Supplementary Figure legend**

**Figure S1. BR inhibits auxin signaling output**

(A) Quantification of the fluorescence signal in the epidermis along the meristem of DR5v2-ntdTomato roots treated with mock, 0.8 nM BL or 3 µM BRZ, for 12 h (n=15,12,22 respectively).

(B) Quantification of the fluorescence signal ($log_{10}$ scale) in the stele of *pIAAmotif-mScarlet-NLS* roots treated with mock, 0. 8nM BL or 3 µM BRZ and 4µM Kyn for 12h (n=15,12,14,11 respectively), as in Figure 1.

**Figure S2. The root meristem of auxin biosynthesis mutant is hypersensitive to BR while epidermal expression of YUC is sufficient to confer high auxin-like phenotype**

(A-C) Phenotype of wild-type and *wei2,7* seedlings after transfer to mock or BL (0.8 nM), for 96h and phenotype of wild type and GL2-YUC5 lines (D-G). (A) Root length (n=15) (B) average cell length versus position from cortex endodermis initial (CEI) (n=8), and (C) whole-seedling phenotype. (D) Root length (n=10) (E) whole-seedling phenotype, (F) meristematic cell length (n=16 and 37 for WT and GL2-YUC5 respectively) and (G) meristem cell number (n=5). The different letters indicate values with statistically significant differences ($p \leq 0.05$). Scale bar = 1cm.

**Figure S3. Dynamics of meristem differentiation in response to hormonal perturbations**

(A) Confocal microscopy images of representative root meristems at 4h-96h after transfer to mock, 0.8 nM BL, 4 μM Kyn or their combination.

(B) Average cell length versus position from CEI (cortex endodermis initial), of seedlings as in (A); see also Figure 2 (n≥8).

(C) Phenotype of 7-day-old seedlings grown as in (A), 96h after transfer to hormonal treatments.

(D) Confocal microscopy images of representative root meristems treated with 100 nM BL or 50 μMu Kyn. Scale bars =100 μm in A and D, scale bar = 1cm in C.

**Figure S4. Simultaneous hormonal perturbation is accompanied by deregulation of gene expression**

(A) DEGs in different root zones at 12 h, 24 h and 48 h. Note that combined BL and Kyn treatment caused concomitant loss of meristematic genes and elevation of differentiation genes by 12h.

(B) Percent of DEGs after 12 h treatment (p≤0.05, and FC≥ 1.5) that are not differentially expressed at 24 h or 48 h of treatment (see also Figure 2).

**Figure S5. The root meristem of BR-deficient mutants is maintained upon a severe block of auxin biosynthesis**

(A) Confocal microscopy images showing the range of meristem phenotype following high-Kyn or combined BL and Kyn treatment. (See Figure 3 and Figure 6)

(B) Seedlings of wild-type, *bri1-116, bri1-GABI* and *cpd,* 10 days after transfer to mock or high Kyn (50 μM). Wild type seedlings were also germinated on BRZ and then subjected to combined BRZ and high Kyn treatment.

(C) 7-day-old wild-type seedlings that were germinated in the presence or absence of 3 μM BRZ before being transferred to high Kyn conditions, for 96 h. In B and C, scale bar = 1cm.

(D,E) Confocal microscopy images and corresponding quantification of *DII-VENUS* 10 days after transfer to mock, 3 μM BRZ, high Kyn (50 μM) or high Kyn+BRZ (n=10), scale bar= 25 μm.

**Figure S6. The root meristem of BR-deficient mutants is maintained upon concomitant moderate perturbation of BR and auxin**

(A,B) Confocal images (A) and seedlings phenotype (B) of wild type, *bri1-116, brl1,brl3 and bri1,brl1,brl3,* 96 h after transfer to mock, BL, Kyn or their combination.

(C) Confocal images of *BIN2/bin2-1* subjected to hormonal treatments as in (A,B). The *BIN2/bin2-1* population segregated to roots, with meristem insensitivity to BL, as indicated by

the wide meristem phenotype (Left root). *BIN2/bin2-1* meristem phenotypes following BL and Kyn treatment for 96h are summarized in a table, see also Figure 6.
Scale bars =100 μm in A and C, scale bar = 1cm in B.

**Figure S7- Phenotypes of transgenic lines with tissue-specific *bin2-1* expression**
(A) Seedling phenotype of wild type and transgenic lines expressing *bin2-1* driven by select tissue-specific promoters: *pWER, pBRN2, pSCR, pSHR and pWOX5*, 96 h after transfer to mock, BL, kyn or their combination (experiment as in figure 5).
(B) Root length of seedlings as in (A) (n≥15).
(C-D) Meristem width (n≥15) (C) and meristematic cortical cell length, measured as distance from CEI (cortex endodermis initial) (D) (n≥8) of wild-type and transgenic lines expressing *bin2-1* in specific root tissues. The different letters indicate values with statistically significant differences between the different lines. The asterisk indicates values that were statistically significantly different from those of the wild type, both according to Tukey HSD ($p \leq 0.05$) (C and D are elaborated experiment as in Figure 5).

### Supplementary Tables

**Table S1- Number of differentially expressed genes (DEGs) in response to mock, 0.8 nM BL, 4μM Kyn and their combinations after 12h, 24h and 48h**

| | Treatment | Up | Down | Total | sum |
|---|---|---|---|---|---|
| 12h | BL | 73 | 42 | 115 | 1461 |
| | Kyn | 163 | 268 | 431 | |
| | BL+Kyn | 448 | 467 | 915 | |
| 24h | BL | 52 | 47 | 99 | 2890 |
| | Kyn | 93 | 160 | 253 | |
| | BL+Kyn | 1486 | 1052 | 2538 | |
| 48h | BL | 81 | 54 | 135 | 3755 |
| | Kyn | 67 | 233 | 300 | |
| | BL+Kyn | 2152 | 1168 | 3320 | |

**Table S2- Primers used in this study**

| Name | Sequence 5' – 3' | Forward (F) /Reverse (R) | Purpose |
|---|---|---|---|
| *pWOX5* | GTCGACGCCAACGTTACAACTTACAAC | F | cloning |
| *pWOX5* | GGTACCGTTCAGATGTAAAGTCCTCAACTG | R | cloning |
| *bin2-1* | TGAAGACATAATGGCTGATGATAAGGAGATGC | F | cloning (GG) |
| *bin2-1* | TGAAGACATCGAACCAGTTCCAGATTGATTCAAGAAGC | R | cloning (GG) |
| *NeonGreen* | TGAAGACATTTCGATGGTCAGCAAGGGCGAGG | F | cloning (GG) |
| *NeonGreen* | TGAAGACATAAGCTTACTTGTAGAGCTCGTCCATGCCC | R | cloning (GG) |
| *Ypet* | TGAAGACATTTCGGTGAGCAAAGGCGAAGAGCTG | F | cloning (GG) |
| *Ypet* | TGAAGACATAAGCTCACTTATAGAGCTCGTTC | R | cloning (GG) |
| *BRI1_CDS* | TGAAGACATAATGAAAACTTTTTCAAGCTTCTTTCTCTC | F | cloning (GG) |
| *BRI1_CDS* | TGAAGACATCGAACCTAATTTTCCTTCAGGAACTTC | R | cloning (GG) |
| *BRI1_mut* | TTGAAGACTTCATCCAACAAAAACCCGTGTACTTTCGAT | F | cloning (GG) |
| *BRI1_mut* | TTGAAGACTTGATGACCAGTCTGGGAGAAGATTCTTGTC | R | cloning (GG) |
| *YUCCA5* | GGTACCATGGAGAACATGTTTAGGCTC | F | cloning |
| *YUCCA5* | GGATCCGGAGCAACTGAAATGCATCTGC | R | cloning |
| *DWF4* | GTTGGCCATTTCTTGGTGAAA | F | Real-time PCR |
| *DWF4* | TGGCGGTGTACGGTTTAAGAT | R | Real-time PCR |
| *CPD* | CCTCCACGATCATGACTCTCG | F | Real-time PCR |
| *CPD* | TTTCATGCTCTTCCTTGAGTTGAG | R | Real-time PCR |
| *DET2* | GGTTATATCCAGGCGAGGTGG | F | Real-time PCR |
| *DET2* | CCATACCGATAACAAACCGCC | R | Real-time PCR |
| *ROT3* | ATTGGCGCGTTCCTCAGAT | F | Real-time PCR |
| *ROT3* | GAACAACCGAGGCCTCAATG | R | Real-time PCR |
| *CYP90D1* | GACGTCTCCAAGACTGTTGC | F | Real-time PCR |
| *CYP90D1* | AGCTCTTCTAAATCTTCTCCTTTCTC | R | Real-time PCR |
| *BROX2* | AACCAGAGGTTGAAGAATATAGAACAGA | F | Real-time PCR |
| *BROX2* | AAATCCACTGCGGTAATTCGTT | R | Real-time PCR |
| *YUCCA9* | CGTAGATGGTCAGAAGCTAGATATCG | F | Real-time PCR |
| *YUCCA9* | AGCCAAGAAGGGACGTTGCT | R | Real-time PCR |
| *At5g15400* | TGCGCTGCCAGATAATACACTATT | F | Real-time PCR |
| *At5g15400* | TGCTGCCCAACATCAGGTT | R | Real-time PCR |
| | | | GG = Golden Gate |
